# A second specificity-determining loop in Class A sortases: Biochemical characterization of natural sequence variation in chimeric SrtA enzymes

**DOI:** 10.1101/2021.03.27.437355

**Authors:** Isabel M. Piper, Sarah A. Struyvenberg, Jordan D. Valgardson, D. Alex Johnson, Melody Gao, Katherine Johnston, Justin E. Svendsen, Hanna M. Kodama, Kelli L. Hvorecny, John M. Antos, Jeanine F. Amacher

## Abstract

Gram-positive bacteria contain sortase enzymes on their cell surfaces that catalyze transpeptidation reactions critical for proper cellular function. *In vitro*, sortases are used in sortase-mediated ligation (SML) reactions for a variety of protein engineering applications. Historically, sortase A from *Staphylococcus aureus* (saSrtA) has been the enzyme of choice for SML reactions. However, the stringent specificity of saSrtA for the sequence motif LPXTG limits its uses. Here, we use principal component analysis to identify a structurally conserved loop with a high degree of variability in all classes of sortases. We investigate the contribution of this β7-β8 loop, located between the catalytic cysteine and arginine residues and immediately adjacent to the target binding cleft, by designing and testing chimeric sortase enzymes. Our chimeras utilize natural sequence variation of Class A sortases from 8 species engineered into the SrtA sequence from *Streptococcus pneumoniae* (spSrtA). While some of our chimeric enzymes mimic the activity and selectivity of the wild-type protein from which the loop sequence is derived (e.g., that of saSrtA), others result in chimeric spSrtA enzymes able to accommodate a range of residues in the final position of the substrate motif (LPXT<u>X</u>). Using mutagenesis, structural, and sequence analyses, we identify three interactions facilitated by β7-β8 loop residues that appear to be broadly characteristic of Class A sortase enzymes. These studies provide the foundation for a deeper understanding of sortase target selectivity and can expand the sortase toolbox for future SML applications.

## Introduction

Sortases are cysteine transpeptidase enzymes that gram-positive bacteria use to covalently attach proteins to their cell wall for various functions, including to assemble pili or display virulence factors (1–3). There are 6 recognized classes of sortase enzymes (classes A-F), with *in vivo* roles ranging from general purpose or “housekeeping” functions (classes A and E), to more specific roles such as the construction of the bacterial pilus (Class C) (1, 4). These enzymes recognize a cell wall sorting signal (CWSS) on the outer membrane of gram-positive bacteria (1, 5). For Class A sortases, the CWSS is the sequence LPXTG (1, 5). Using previously published numbering (L=P4, P=P3, X=P2, T=P1, and G=P1’), P4, P3 and/or P1’ of this motif vary amongst different classes (5). Following target recognition, a His-Cys-Arg catalytic triad facilitates a ligation reaction whereby the CWSS is cleaved between threonine and glycine residues, followed by resolution of an acyl-enzyme intermediate via nucleophilic attack by an incoming amino group that results in formation of a new peptide bond (1, 3, 5, 6).

The ability to cleave a signal sequence and subsequently ligate a second component (typically a protein or synthetic peptide derivative) via a covalent bond make sortases an attractive tool for protein engineering efforts, commonly called sortase-mediated ligation (SML) or *sortagging* (3). Sortase A from *Staphylococcus aureus* (saSrtA) was the first of these enzymes discovered and continues to see widespread use for *in vitro* SML experiments (**Fig. 1A-B**) (1, 7). Recent years have seen continuous improvements in SML technology, including strategies for limiting the reversibility of the ligation reaction, and the development of saSrtA variants with dramatically improved catalytic efficiency (3, 8). However, in the majority of cases SML remains restricted to the pentapeptide LPXTG motif, which limits its utility as a protein engineering tool (9, 10). A relaxed recognition motif could potentially allow scientists to target a larger number of protein targets that do not contain an endogenous LPXTG sequence (11).

**Fig. 1.**
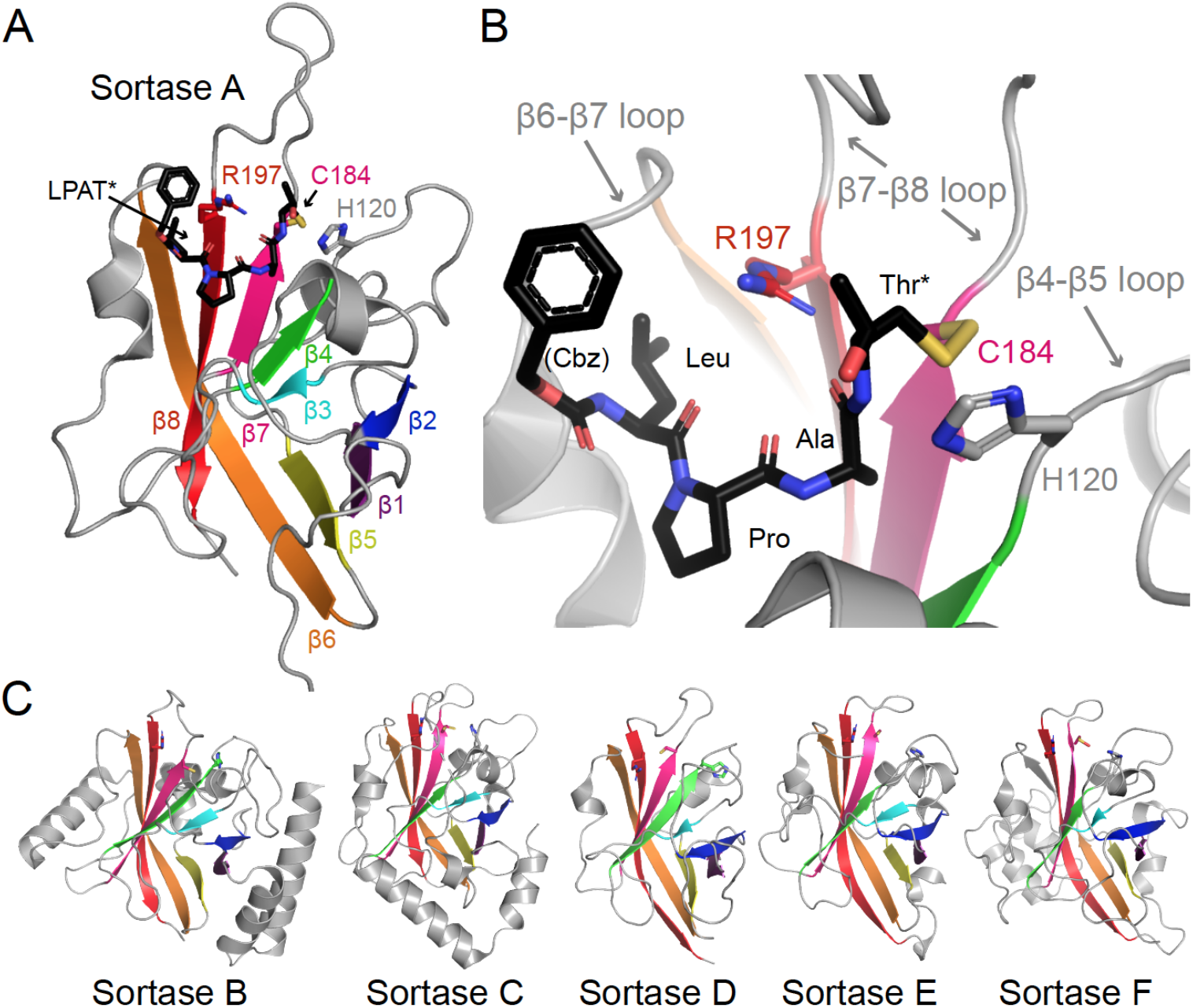
The sortase-fold is conserved in all classes of bacterial sortases. (A) The peptide-bound structure of *S. aureus* SrtA (saSrtA) is shown in cartoon representation, with β-strands colored and labeled (PDB ID 2KID) (16). The side chains of the catalytic residues (H120, C184, and R197) are shown as sticks, colored by heteroatom (O=red, N=blue, S=yellow), and labeled. The peptide analog, Cbz-LPAT*, where Cbz is a carbobenzyloxy protecting group and T* is (2*R*,3*S*)-3-amino-4-mercapto-2-butanol, is shown as black sticks and colored by heteroatom (16). A zoomed-in version of the active site is shown in (B), with features indicated as in (A). The variable loops are labeled and indicated by gray arrows. (C) The overall sortase-fold is well conserved in proteins of different classes. Here, structures for Class B (PDB ID 1NG5), Class C (3O0P), Class D (2LN7), Class E (5CUW), and Class F (5UUS) sortases are in cartoon, with conserved β-strands colored as in (A), highlighting the 8-stranded sortase-fold.

Previous mutagenesis and structural studies of various sortases provide a wealth of knowledge about initial ligand recognition and subsequent cleavage (thioesterification), as well as nucleophile recognition and mechanistic details of peptide ligation (transpeptidation) (1, 2, 9). Specifically, the catalytic residues of all native sortases identified to date are (using saSrtA numbering unless specified otherwise): His120 (general acid/base), Cys184 (nucleophile, acyl-enzyme intermediate), and Arg197 (transition state stabilization) (**Fig. 1B**) (1, 9). Additionally, directed evolution studies have identified mutations (P94R/D160N/D165A/K190E/K196T) that are able to boost the catalytic efficiency of saSrtA by 120-fold (8). Of these 5 mutations, several are located in two of the three structurally conserved loops in Class A sortases: those between the β4, β5 strands (β4-β5 loop), the β6, β7 strands (β6-β7 loop, where D165A occurs), and the β7, β8 strands (β7-β8 loop, where K190E and K196T are located). Notably, while the increase in enzyme activity afforded by these mutations included a 3.6-fold increase in k_cat_, the effect was dominated by a 33-fold decrease in K_M_, suggesting these loop residues may be important in CWSS recognition (8).

Additional evidence for the role of loop residues has been obtained from more targeted directed evolution and mutagenesis studies. For example, it has been demonstrated that the β6-β7 loop of saSrtA directly confers specificity at P4 of the recognition motif (<u>L</u>PXTG), and residues other than leucine (L) can be accommodated using sortases with mutations in the β6-β7 loop. (12–14). Indeed, substitution of the β6-β7 loop residues from saSrtB into the saSrtA enzyme alters substrate recognition to that of a sortase B protein (NPQTN) (15). Turning to the β7-β8 loop, the NMR structure of saSrtA covalently bound to a modified LPAT* peptide mimetic revealed a non-covalent interaction between W194 in saSrtA and the Thr residue in P1 (LPX<u>T</u>G) (16, 17). Mutation of W194 in saSrtA decreased the reaction rate, although it was not essential to catalysis (17). Taken together, these past studies reveal that sequence variation within sortase loops directly affects both activity and selectivity for target ligands. Furthermore, conservation of the closed eight-stranded β-barrel core in all sortase A-F structures that have been reported to date suggests that these principles may apply to non-Class A sortases as well. (**Figs. 1A, C**) (2).

In this work we specifically look at natural sequence variation in the β7-β8 loop of Class A sortases, using *Streptococcus pneumoniae* SrtA as a model system. The β7-β8 loop was initially identified using principal component analysis as a region of notable variability in Class A (and other) sortases. We find that the β7-β8 loop sequence dramatically affects both overall enzyme activity, as well as selectivity at P1’ of the CWSS. Our data is consistent with a recent publication that investigated the grafting of β7-β8 loop sequences from saSrtA and *Bacillus anthracis* SrtA into *Streptococcus pyogenes* SrtA (18). This work also suggested that W194 (saSrtA numbering) may play a role in the substrate recognition of the reported chimeras (18). Here, we have profiled the substrate preferences of over a dozen loop chimeras and single- or double-mutants targeting the β7-β8 loop. While we also observe a role for W194 in substrate recognition, our data suggests that it is unique to saSrtA and not broadly applicable to describe β7-β8 loop-mediated Class A sortase function. Indeed, the combination of functional enzyme assays and analysis of reported sortase structures in the present work suggests three different β7-β8 loop-mediated interactions that affect selectivity and activity.

## Results

### Principal component analysis (PCA) of bacterial sortases

In order to gain a better understanding of global sequence patterns in the sortase superfamily, we used principal component analysis (PCA) to group and analyze 39,188 sortase sequences from all classes (Experimental Procedures). Briefly, we downloaded all sequences annotated as “sortase” from UniProt and aligned then by MAFFT, followed by PCA (19, 20). The amino acids in each sequence were then classified by 5 parameters: hydrophobicity, disorder propensity, molecular weight, charge, and occupancy (defined as a binary value, where 1 = amino acid and 0 = insertion or deletion (indel) at this position) (21, 22). PCA was then performed on the resulting 39,188*5-dimensional data. This data was projected onto 3 principal components where 42.7% of the total variance is described (**Figs. S1A**). Hierarchical clustering of the sortase superfamily was achieved by using the first two components of the singular value decomposition (SVD) matrix. The projected points were then partitioned into 2 clusters by Gaussian mixture modeling. This process is performed recursively on the dataset until each cluster reaches a minimum size or the Gaussian mixture modeling process fails to identify two distinct gaussians (23). The resulting tree from this process can accurately distinguish the known sortase classes (**Fig. 2A**). We also plotted our PCA using the top three principle components (**Fig. S1B**). For visualization, we ran PCA on a subset of the data, including 9,427 sequences that were filtered for low numbers indels and manually verified (**Fig. 2B**).

**Fig. 2.**
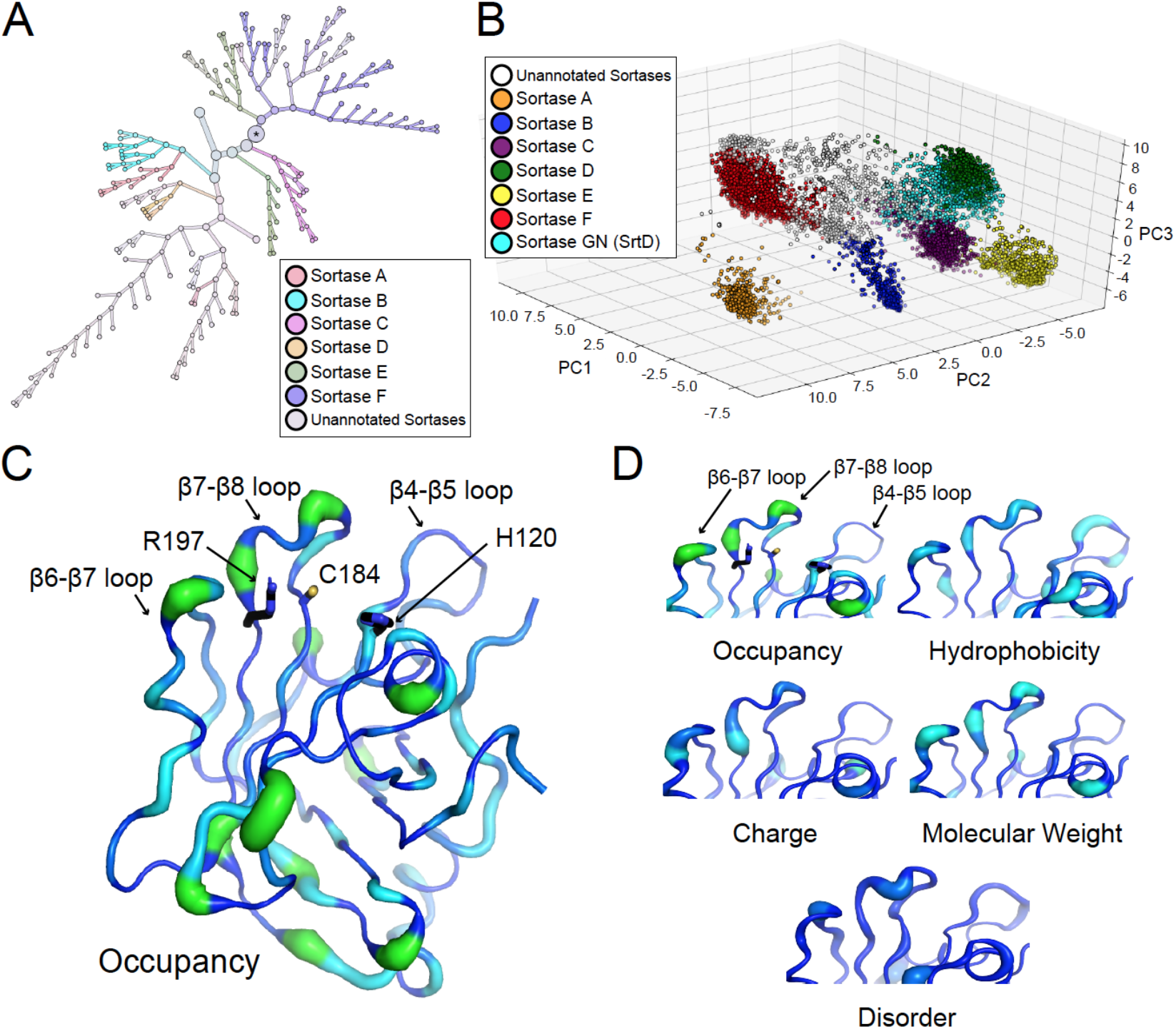
Principal component analysis (PCA) of sortase superfamily reveals sequence variability in structurally-conserved loops. (A) Hierarchical clustering of the sortase superfamily by gaussian mixture model unsupervised classification on the first two components of the singular value decomposition (SVD) matrix distinguishes the known classes of sortases (23). (B) Visualization of a subset of 9,427 of the sortase sequences, scored and filtered with respect to a low number of insertions or deletions (indels) and plotted using principle components 1-3. The sequence is colored by which class of sortase it is annotated as by UniProt, when available. An equivalent plot of all 39,188 sortase sequences is in **Fig. S1C**. (C-D) The 5 characteristics assigned numerical values in the PCA are visualized by width (from 0 to 1) and color (where darker is a value closer to 0 and lighter indicates a value closer to 1 using PyMOL. The *Streptococcus pyogenes* SrtA structure (PDB ID 3FN5) is used as the model to show variance in a “typical Class A sortase.” The catalytic residues (H142, C208, and R216) are shown as side chain sticks and colored by heteroatom. The 3 structurally conserved loops that are discussed in this work are labeled. We focused on variance near the active site here, but notably, there is also a relatively large degree of variance on the other side of the protein (also **Fig. S1D**).

This analysis verified previous classifications of sortases based on sequence alignment, network, and phylogenetic tree analyses (4, 24, 25). For example, principal component 1 (PC1) separates the sortase F proteins from the rest of the superfamily and PC2 captures the separation between sortase B and the other sortase families, as well as sortase E and sortase A. These analyses allowed us to identify the regions of highest variability within each class based on the parameters defined above. We plotted our data onto previously determined sortase A structures by linearizing the distance from the centroid for each position in the multiple sequence alignment (**Fig. S1C**). Unsurprisingly, we found that secondary structure elements are highly conserved, including the “sortase fold”β-barrel core and class-specific a-helices (**Figs. 2C-D and S1D**). Additionally, PCA revealed that the highest degree of variability occurs in structurally conserved loops adjacent to the substrate recognition pocket (**Figs. 2C-D and S1D**).

Given that the β6-β7 loop has been shown to be intimately involved in sortase substrate recognition, we were intrigued that PCA revealed similar levels of variability in the β4-β5 and β7-β8 loops (15). In the case of β7-β8, we were also motivated by previously reported mutations in the β7-β8 loop of saSrtA that have been shown to dramatically modulate sortase reaction rates (8, 17, 26). Therefore, we sought to further explore how the β7-β8 loop affects the activity and substrate specificity of a sortase with narrow substrate tolerance (saSrtA) versus one that is more promiscuous (spSrtA).

### Loop-swapped β7-β8 variants reveal differences in position P1’ selectivity for S. aureus and S. pneumoniae SrtA enzymes

In our previous work, we found that while saSrtA is specific for a Gly residue at P1’ (LPXT<u>G</u>) of the substrate motif, SrtA from *Streptococcus pneumoniae* (spSrtA) recognizes over 10 of the 20 amino acids at this position in a 24-hr end point assay (27). To determine whether the β7-β8 loop played a role in these differing substrate preferences, we began by engineering two loop-swapped variants: saSrtA_pneumoniae_ (which contains the β7-β8 loop residues from spSrtA (<u>C</u>EDLAATE<u>R</u>, where the catalytic cysteine and arginine are underlined), and spSrtA_aureus_ (with β7-β8 residues <u>C</u>DDYNEKTGVWEK<u>R</u> from saSrtA). Notably, the length of the saSrtA β7-β8 loop contains an additional five residues, as compared to the spSrtA β7-β8 loop. The saSrtA loop also uniquely contains W194, which is known to directly contact the P1 threonine of the LPXTG motif (16). In addition, while both loops are predicted to have an overall net negative charge at physiological pH, the saSrtA loop contains two positively charged lysine residues that are not present in spSrtA. Both chimeric sortases were expressed and purified from *E. coli*, and were isolated as soluble, monomeric enzymes as described previously and in the Experimental Procedures (**Fig. S2A**) (27). The secondary structure content for all variants was consistent with the respective wild-type protein, as measured by circular dichroism and described in the Experimental Procedures (**Fig. S2B**).

To monitor enzymatic activity and selectivity of the saSrtA_pneumoniae_, spSrtA_aureus_, and their wild-type counterpart proteins, we utilized well-established FRET quencher probes consisting of different substrate motifs flanked by a 2-aminobenzoyl fluorophore (Abz) and a 2,4-dinitrophenyl quencher (Dnp) (17, 28, 29). Probes containing three substrate variants were initially prepared (Abz-LPAT**A**G-K(Dnp), Abz-LPAT**G**G-K(Dnp), Abz-LPAT**S**G-K(Dnp), varying only at P1’ in **bold**) and used to test the relative activity of our wild-type and chimeric enzymes (**Table S2**). For simplicity, we have hereafter omitted the Abz, K(Dnp), and C-terminal glycine from peptide descriptions. For comparing enzyme activity, a standard 2 h reaction time was utilized, and an excess of H_2_NOH was included to resolve the acyl enzyme intermediates. For consistency, all reactions were also conducted in the presence of Ca^2+^, which is a required co-factor for saSrtA. Reaction endpoint (indicated by the increase in Abz fluorescence) for all enzyme/substrate pairings was then expressed relative to averaged benchmark reactions of wild-type saSrtA with the standard LPAT**G** substrate (**Figs. 3A, S3A**). This benchmark reaction was consistently found to give ~84% conversion to the expected transacylation products when independently monitored via RP-HPLC (**Fig. S3B**).

**Fig. 3.**
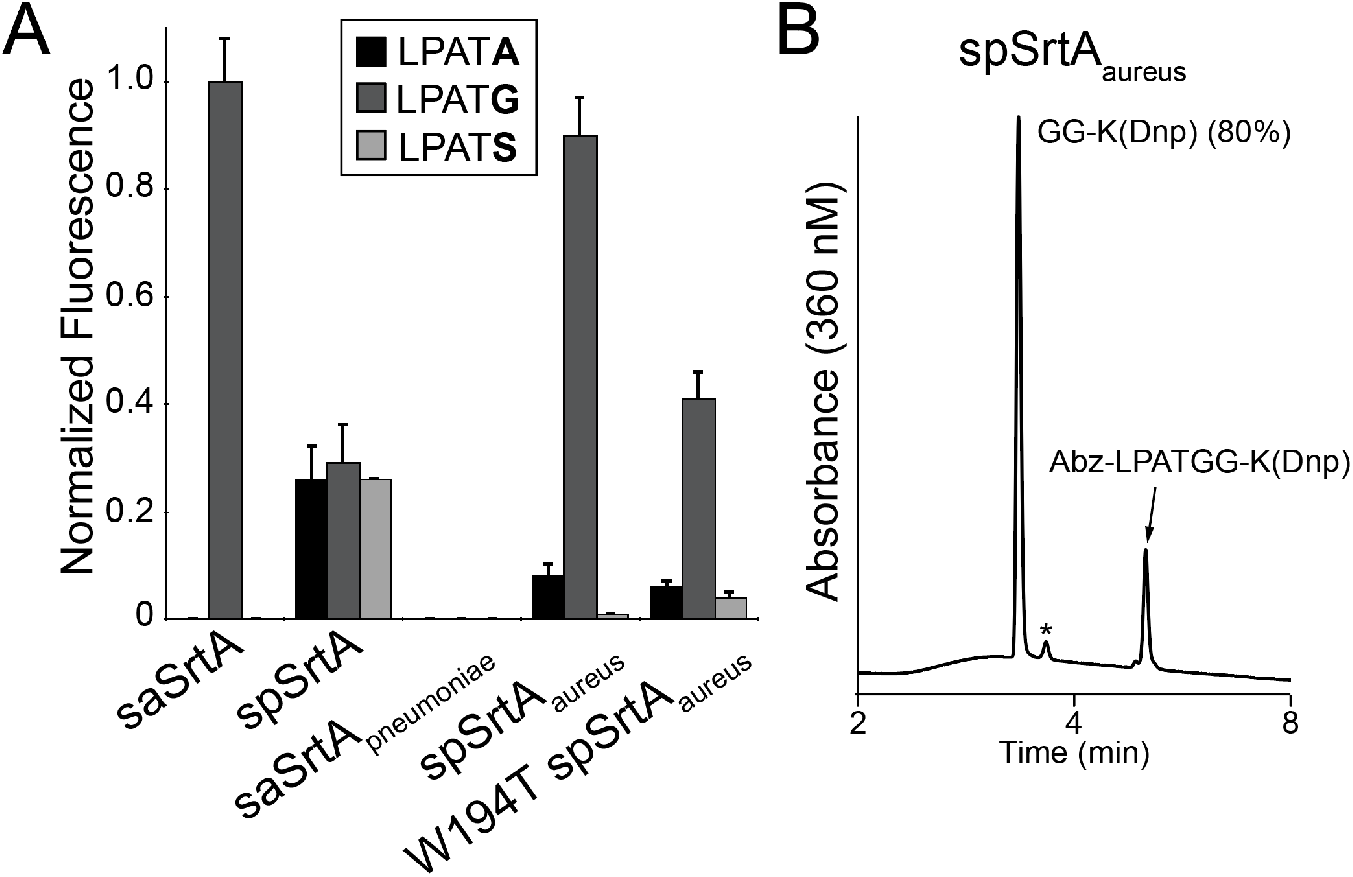
Interchanging β7-β8 loops in class A sortases modulates substrate selectivity and activity for target sequences that vary at position P1’ of the canonical LPXTG motif. (A) Comparison of substrate selectivity for wild-type saSrt and spSrtA proteins, as well as β7-β8 loop chimeras saSrtA_pneumoniae_, spSrtA_aureus_, and W194T spSrtA_aureus_. Substrate cleavage was monitored via an increase in fluorescence at 420 nm from reactions of the fluorophore-quencher probes Abz-LPATGG-K(Dnp), Abz-LPATAG-K(Dnp), and Abz-LPATSG-K(Dnp) (represented as LPAT**G**, LPAT**A**, and LPAT**S**) in the presence of excess hydroxylamine. Bar graphs represent normalized fluorescence (± standard deviation) from triplicate experiments at the 2 h reaction timepoint, as compared to saSrtA and the peptide LPAT**G** (raw values are in **Table S1**). (B) Representative HPLC chromatogram for the reaction of Abz-LPATGG-K(Dnp) and H_2_NOH in the presence of spSrtA_aureus_. This reaction was conducted in the presence of Ca^2+^. Selective cleavage between the threonine (T) and glycine (**G**) residues was observed, with an overall conversion of 80% (* = Abz-LPAT-NHOH reaction product. Low peak intensity is due to the weak absorbance of Abz at 360 nm). Also see **Fig. S3B-F**.

Based on our previous results, we predicted that spSrtA would show activity for all three peptides, while saSrtA would be selective for LPAT**G** (27). Consistent with this prediction, our results confirmed that spSrtA was equally capable of processing all three substrates, whereas saSrtA was restricted to LPAT**G** (**Fig. 3A**). Our assay also revealed a marked reduction in spSrtA activity versus saSrtA, which was not captured in our previous study, likely due to the extended reaction time (24 h) used in that work (27). With respect to the chimeric enzymes, our results clearly showed that the sequence of the β7-β8 loop was a major determinant of activity and specificity. Specifically, the saSrtA_pneumoniae_ protein was completely inactive while spSrtA_aureus_ functionally mimicked the narrow substrate preference of the wild-type saSrtA enzyme (**Fig. 3A**). This result is consistent with recently published data (18). To verify that our sortases were cleaving substrates at the expected site, reactions exhibiting a normalized fluorescence value of 0.2 were independently monitored by RP-HPLC and LC-MS, which confirmed cleavage between P1 and P1’ (**Figs. 3B, S3B-F**). Notably, reactions for HPLC and LC-MS characterization were conducted in the presence and absence of Ca^2+^, which demonstrated that this cofactor was not required for the activity of spSrtA and spSrtA_aureus_.

Continuing on with the SpSrtA_aureus_ chimera, we next wanted to determine if the Trp residue derived from the saSrtA loop played a significant role in enzyme activity. In wild-type saSrtA, the W194 residue (using saSrtA numbering) is known to affect enzyme activity of saSrtA via direct interactions with the threonine of the LPXTG motif (16, 17). We therefore expressed and purified the corresponding “W194T” mutant of spSrtA_aureus_ and tested this variant with A-, G-, and S-containing peptides in our assay. Indeed, our W194T spSrtA_aureus_ protein exhibited a 59% reduction in reaction progress for LPAT**G**, while retaining its selectivity for Gly-containing peptides (**Fig. 3A, Table S1**). This result suggests that W194 likely interacts with the peptide substrate in a similar manner as previously described, despite the spSrtA scaffold (16).

### Variability in position P1’ selectivity and transpeptidase activity in S. pneumoniae SrtA β7-β8 variants

In addition to the profound shift in substrate scope observed for spSrtA_aureus_, we were also intrigued that the overall reactivity of this chimera for LPAT**G** was comparable to that seen with wild-type saSrtA. This stood in sharp contrast to the reaction of LPAT**G** with wild-type spSrtA, where reaction progress was nearly two-thirds lower within the 2 h reaction time of our assay (**Fig. 3A**). Based on this, we wondered if similar gains in reactivity for substrates other than LPAT**G** could be achieved by substituting in residues from additional SrtA proteins (27, 28). To test this, we created an additional 6 spSrtA variants containing loop residues from SrtA proteins that we had evaluated previously (27). These chimeras included the β7-β8 loop residues from *Bacillus anthracis* (spSrtA_anthracis_), *Enterococcus faecalis* (spSrtA_faecalis_), *Lactococcus lactis* (spSrtA_lactis_), *Listeria monocytogenes* (spSrtA_monocytogenes_), *Streptococcus oralis* (spSrtA_oralis_), and *Streptococcus suis* (spSrtA_suis_) (**Fig. 4A**) (27). To avoid confusion in the numbering of loops with variable lengths, we will hereafter refer to the N-terminal positions of each β7-β8 loop by numbering with respect to the catalytic Cys (β7-β8^+1^, β7-β8^+2^, etc.) that precedes the loop, whereas the C-terminal loop residue will be numbered relative to the catalytic Arg (β7-β8^−1^) (**Fig. 4A**).

**Fig. 4.**
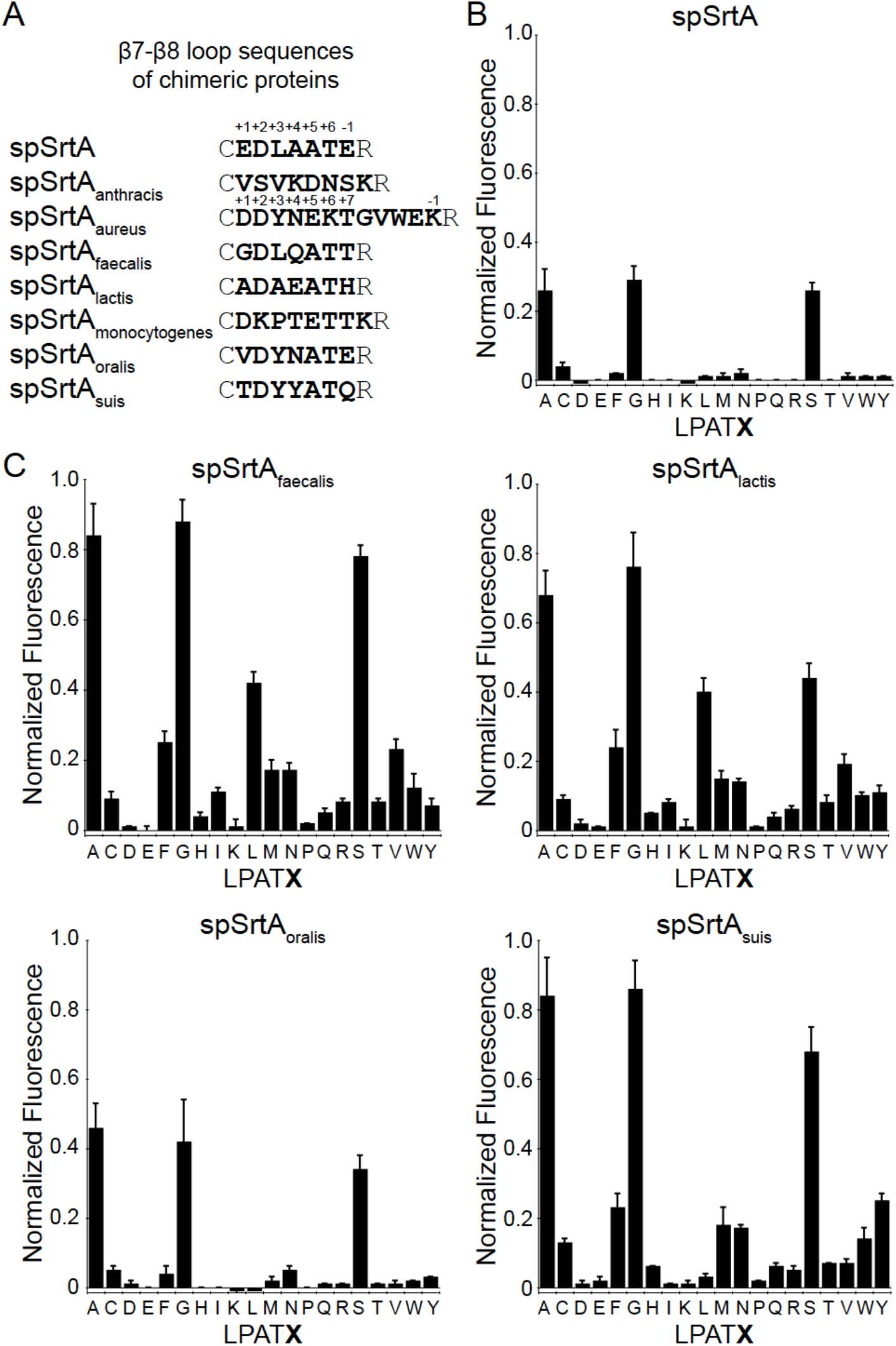
The sequence of the β7-β8 loop dramatically affects selectivity and activity for spSrtA. (A) The β7-β8 loop sequences of the chimeric proteins used are listed, with representative numbering for residues in the β7-β8 loop labeled for spSrtA and spSrtA_aureus_. (B-C) Substrate selectivity profiles for wild-type spSrtA (B) and chimeric spSrtA variants (C). Substrate cleavage monitored via an increase in fluorescence at 420 nm from reactions of fluorophore-quencher probes with the generic structure Abz-LPATXG-K(Dnp) (LPAT**X**) in the presence of hydroxylamine. Bar graphs represent mean normalized fluorescence (± standard deviation) from at least three independent experiments.

The 6 chimeric proteins were expressed and purified using the same protocol as spSrtA_aureus_, and as described in the Experimental Procedures. The purity of all proteins was validated by SDS-PAGE, and size exclusion chromatography was consistent with the isolated proteins being predominantly monomeric (**Fig. S2A**). In addition, all variants contained similar secondary structure content as compared to the respective wild-type protein, as measured by circular dichroism (**Fig. S4A**). With the new chimeras in hand, we conducted an initial evaluation of relative activity using the LPAT**G**, LPAT**A**, and LPAT**S** substrates described above. While the majority of constructs exhibited significant reactivity across all three substrates, the spSrtA_anthracis_ and spSrtA_monocytogenes_ proved to be inactive (**Fig. S4B**). For the remaining enzymes, the spSrtA_oralis_ protein behaved similarly to wild-type spSrtA, while spSrtA_faecalis_, spSrtA_lactis_, and spSrtA_suis_ showed improved performance for A-, G-, and S-containing substrates (**Figs. 4B-C, S4B**). This was particularly interesting in the case of spSrtA_faecalis_ given that the wild-type SrtA enzyme from *E. faecalis* was previously shown to have poor reactivity for the same test substrates despite the use of higher enzyme loading and considerably longer reaction times (27).

Based on initial experiments with the A-, G-, and S-containing peptides, we next wanted to expand our peptide pool in order to assess the relative reactivity of our active chimeric spSrtA variants for peptides containing all 20 amino acids at P1’. For comparison, a similar substrate profile was generated for wild-type spSrtA. As shown in **Fig. 4B**, within the 2 h time frame of our assay, the wild-type protein was rather selective in its substrate recognition, with reactivity limited to A-, G-, and S-containing peptides (**Table S1**). We note here that this somewhat limited substrate scope appears to differ from the more promiscuous behavior reported previously for spSrtA. We attribute this to the fact that longer reaction times (24 h) and higher enzyme loadings (5-fold higher than the loading used here) were utilized in this earlier work (27). Similar to wild-type spSrtA, the spSrtA_oralis_ was limited to A-, G-, and S-containing peptides, albeit with slightly elevated reactivity in the case of LPAT**A** and LPAT**G**. Finally, we were intrigued to find that our spSrtA_faecalis_, spSrtA_lactis_, and spSrtA_suis_ proteins all show increased promiscuity for a variety of amino acids at P1’ in our assay (**Fig. 4C, Table S1**).

Overall, the spSrtA_faecalis_, spSrtA_lactis_, and spSrtA_suis_ proteins showed the largest increase in activity and promiscuity for this library of peptides. The spSrtA_faecalis_ and spSrtA_lactis_ proteins each recognized 15 of the 20 amino acids at P1’ with normalized fluorescence values of ≥ 0.05, while spSrtA_suis_ recognized 14 of the 20 (**Table S1**). We chose 0.05 as a cut-off value in order to compare with the peptide activities of the spSrtA protein, which shows normalized fluorescence values of −0.02 to 0.02 for all non-G-, S-, or A-containing peptides, with the exception of LPAT**C** (at 0.04 ±0.01) (**Table S1**). Furthermore, spSrtA_faecalis_ and spSrtA_suis_ exhibited ~3-fold higher reaction progress for the G-, S-, and A-containing peptides, as compared to spSrtA (**Table S1**).

As verification of the results of our fluorescence assay, we also characterized a subset of enzyme/substrate combinations using RP-HPLC and LC-MS. Focusing on spSrtA_faecalis_ we repeated reactions that exhibited normalized fluorescence values of ≥ 0.1 (LPAT**X**, **X** = A, F, G, I, L, M, N, S, V, W, Y) (**Fig. S4C**). Reactions were conducted in the absence of Ca^2+^ to confirm that this cofactor was not required for activity. Successful substrate cleavage was observed in all cases, ranging from a high of 78% conversion in the case of LPAT**G**, to only 6% conversion in the case of LPAT**W** over 2 hours at temperature (**Fig. S4C**). Notably, the trends in relative substrate preferences observed by HPLC were consistent with those found in our original fluorescence assay (**Fig. S5**). Additionally, while LC-MS characterization confirmed that substrate cleavage was occurring between the P1 and P1’ of all sequences, certain substrates (LPAT**X**, **X** = W,F,L) containing bulky hydrophobic residues also produced alternate products arising from cleavage on the C-terminal side of P1’. In the case of LPAT**L**, this alternate cleavage product was actually the major species obtained following reaction with spSrtA_faecalis_. We note here that this capacity for alternate cleavage has been reported previously for wild-type spSrtA, and thus appears to be maintained in the spSrtA_faecalis_ chimera (27).

### Variability in position P1’ selectivity and ligase activity in S. pneumoniae SrtA β7-β8 variants

As a final assessment of the reactivity of the spSrtA_faecalis_ chimera, we next evaluated its ability to ligate amino acid nucleophiles in place of the H_2_NOH that was utilized in our fluorescence assay. For a series of test substrates (LPAT**X**, **X** = A,S,V), spSrtA_faecalis_ was able to successfully ligate the corresponding free amino acid carboxamides (**X**-NH_2_ = A-NH_2_, S-NH_2_, V-NH_2_) with very good efficiency (**Fig. 5**). As expected from our fluorescence assay results, reaction progress with LPAT**V** was slower than that observed for LPAT**A** and LPAT**S**. Specifically, reactions with LPAT**V** required 8 h at room temperature to consume 85% of the initial peptide substrate, whereas reactions with LPAT**A**/**S** exhibited >95% substrate conversion within 3 h. Importantly, the desired LPAT**X**-NH_2_ species was the major ligation product in all reactions as determined by LC-MS (**Fig. 5**, **Table S2)**. Trace levels of substrate hydrolysis were also observed via LC-MS, however the ratio of successful ligation to hydrolysis was 15:1 or better as estimated from mass spectral peak intensities. In reactions involving LPAT**V**, we also detected low levels of substrate cleavage on the C-terminal side of the P1’ valine residue. The extent of this alternate cleavage pathway was minimal, accounting for only ~4% of the substrate cleavage events based on comparisons of HPLC peak areas for G-K(Dnp) and **V**G-K(Dnp) (**Fig. 5**). Interestingly, LC-MS characterization of these same reactions involving LPAT**V**, **V**-NH_2_, and spSrtA_faecalis_ failed to show clear evidence for the formation of ligation or hydrolysis products derived from the alternate cleavage pathway, potentially due to their low levels in solution.

**Fig. 5.**
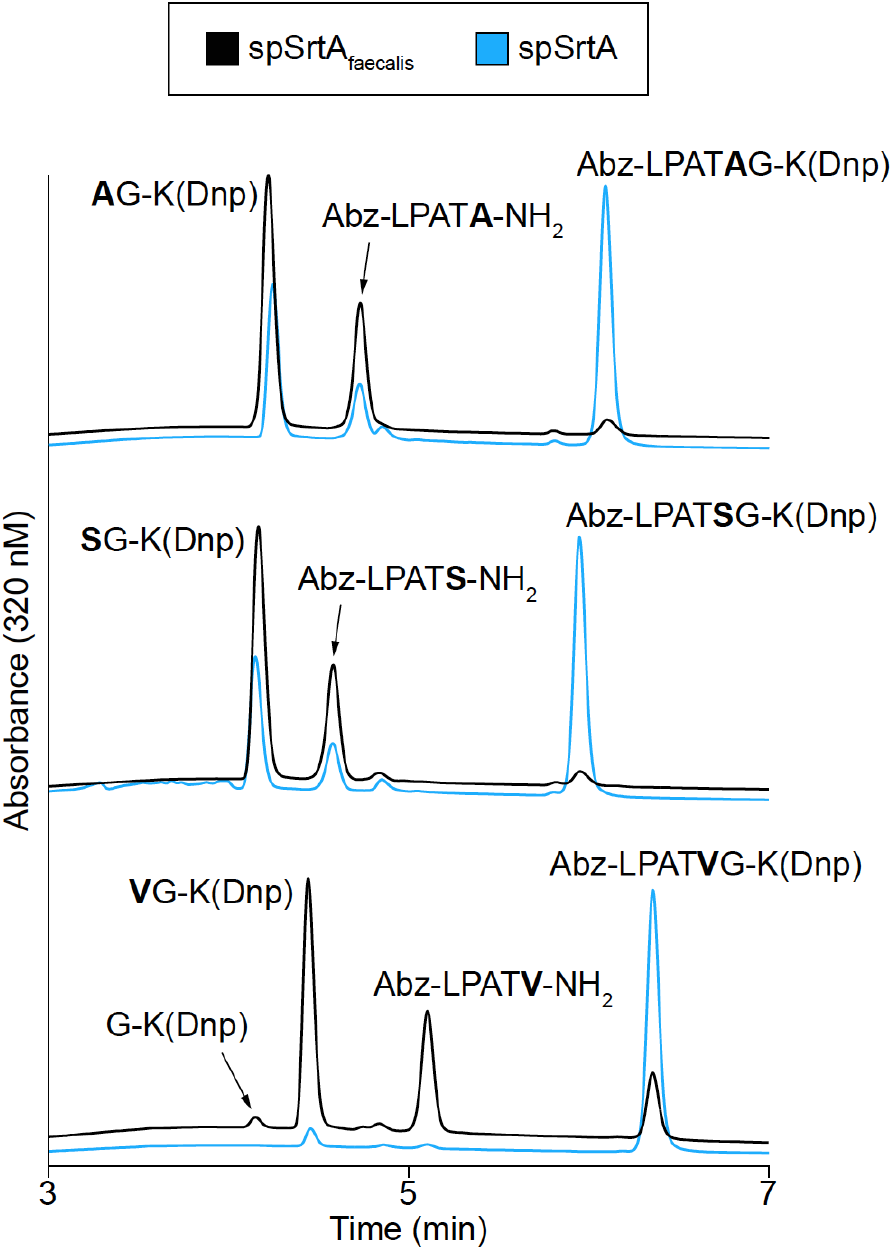
spSrtA_faecalis_ outperforms wild-type spSrtA in model amino acid ligation reactions. HPLC chromatograms (320 nm) for model ligations between Abz-LPATXG-K(Dnp) and excess X-NH_2_ nucleophiles catalyzed by spSrtA_faecalis_ (black curves) or wild-type spSrtA (blue curves). Ligations were conducted in the absence of Ca^2+^. Chromatograms for LPAT**A**/**S** represent the 3 h reaction timepoint, and chromatograms for LPAT**V** correspond to the 8 h timepoint. All peaks identities were confirmed via LC-MS (**Table S2**).

For comparison, we also performed the same set of test ligations with wild-type spSrtA. In all cases, reaction progress was significantly reduced as compared to spSrtA_faecalis_ (**Fig. 5**). In particular, spSrtA exhibited minimal product formation with the LPAT**V** system, representing a 10-fold reduction in reaction progress relative to spSrtA_faecalis_ for this atypical sortase substrate motif. Building from this result, an initial attempt to utilize the LPAT**V** sequence as a handle for site-specific protein modification was made by installing this motif at the C-terminus of a full-size protein target. However, this protein substrate proved to be unreactive in the presence of both spSrtA_faecalis_ and wild type spSrtA (data not shown).

### Stereochemical basis of β7-β8 variant selectivity and activity

In order to gain a stereochemical understanding of our biochemical results, we first analyzed available structures of Class A sortases in the Protein Data Bank. To our knowledge, the 3D structure of an active, monomeric form of spSrtA has yet to be reported. Available crystal structures of the domain-swapped dimer show that the β7-β8 loop is located at, and participates in, the dimer interface (PDB codes 4O8L, 4O8T, and 5DV0). Therefore, we chose to broaden our search to non-spSrtA structures, and in doing so we identified 3 putative β7-β8 loop-mediated interactions in Class A sortases:

#### (1) A Thr-mediated intra-loop hydrogen bond

We initially observed an intra-loop hydrogen bond formed in the β7-β8 loops of several Class A sortases (**Fig. 6A**). This Thr-mediated hydrogen bond is evident in SrtA proteins from *S. aureus* (PDB code 2KID), *S. pyogenes* (3FN5), *S. mutans* (4TQX), and *L. monocytogenes* SrtA (5HU4), among others not shown (**Fig. 6A**). We predict that in spSrtA, this hydrogen bond will be formed between the side chains of residues β7-β8^+2^ D209 and β7-β8^+6^ T213. Indeed, we see this interaction in a homology model of the monomeric spSrtA protein generated using SwissModel (**Fig. S6A**). For this model, the *Streptococcus pyogenes* SrtA structure (PDB code 3FN5) was used as a template because the crystallized form of this enzyme (*S. pyogenes* SrtA residues S81-T249) has 63% sequence identity with spSrtA (30–32). An alignment of our spSrtA model with 3FN5 revealed an overall RMSD of 0.083 Å over 567 main chain atoms. We further validated our model using structural alignments with a monomer extracted from the domain-swapped dimer structure (RMSD of 0.603 Å over 483 main chain atoms), as well as other SrtA structures from *Streptococcus* species, including those from *Streptococcus agalactiae* and *Streptococcus mutans* (PDB codes 3RCC (RMSD of 0.773 Å over 384 main chain atoms) and 4TQX (RMSD of 0.456 Å over 530 main chain atoms), respectively) (**Fig. S6B-C**).

**Fig. 6.**
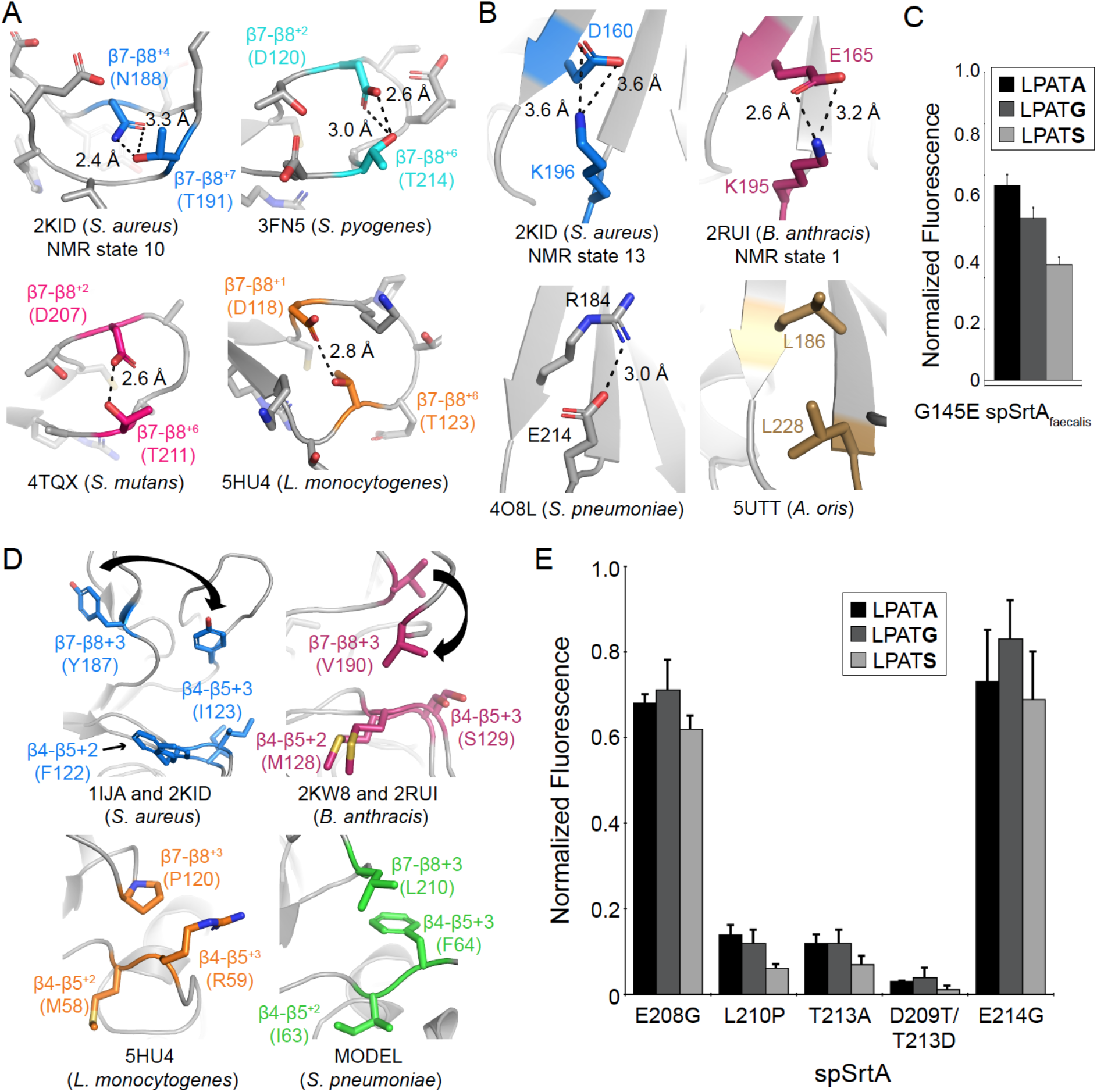
Residues in the β7-β8 loop participate in interactions that affect enzyme activity and selectivity, based on structural analyses and mutagenesis. (A-B, D) All SrtA structures are in gray ribbon. The β7-β8 loop side chains are all in stick representation and colored by heteroatom. Residues that participate in hydrogen-bonds have non-gray carbons and are labeled. The hydrogen bonds are shown with a black dashed line, with measurements indicated. For NMR structures (PDB IDs 2KID and 2RUI), the state that was used for the image is labeled. While not all NMR states contained the interaction indicated, in both cases, there were several states that revealed measurements consistent with a non-covalent interaction. Any side chain sticks are colored as described below colored by heteroatom (O=red, N=blue). (A) *β7-β8 intra-loop hydrogen bond*.The structures in this figure are: *S. aureus* SrtA (2KID, blue carbons), *S. pyogenes* SrtA (3FN5, cyan carbons), *S. mutans* SrtA (4TQX, pink carbons), and *L. monocytogenes* SrtA (5HU4, orange carbons). (B) *Interaction between the β7-β8 loop and β6 strand*. The structures in this figure are: *S. aureus* SrtA (PDB ID 2KID, blue carbons), *B. anthracis* SrtA (2RUI, dark pink carbons), *S. pneumoniae* SrtA (4O8L, gray carbons), and *A. oris* SrtA (5UTT, gold carbons). (C) The G145E mutation in spSrtA_faecalis_ reduces activity for a representative set of target peptides (fluorophore-quencher substrate assay conditions identical to those described in **Fig. 3**). Bar graphs represent normalized fluorescence values (± standard deviation) from three independent experiments. (D) *Interaction between the β7-β8 and β4-β5 loops*. The structures in this figure are: *S. aureus* (unbound: 1IJA and bound: 2KID, blue carbons), *B. anthracis* (unbound: 2KW8 and bound: 2RUI, dark pink carbons), *L. monocytogenes* (5HU4, orange carbons), and *Streptococcus pneumoniae* (homology model (**Fig. S6**), green carbons). The arrows indicate residue movement from the unbound to bound structures. (E) Mutagenesis activity data for spSrtA and G-, S-, and A-containing peptides. Conditions identical to (C).

We hypothesize that this β7-β8 intra-loop hydrogen bond is important for reducing flexibility in the β7-β8 loop, which may be an important characteristic of certain sortases prior to substrate recognition. This is apparent in saSrtA (PDB code 2KID), where NMR structures reveal that the β7-β8 loop is ordered in both the unbound and bound states (**Fig. S7A**). In contrast, previously published NMR structures of the *B. anthracis* SrtA (baSrtA) protein (PDB codes 2KW8 and 2RUI), which does not contain a β7-β8 loop Thr, point to a distinct mechanism. These structures show that a unique N-terminal appendage in baSrtA regulates active site accessibility and that the β7-β8 loop transitions from a disordered-to-ordered state upon substrate binding (**Fig. S7B**) (33). In the ordered/bound state, there are β7-β8 intra-loop hydrogen bonds, specifically between two Ser residues with the peptide backbone and/or D192, but overall, these interactions appear to be distinct from the Thr-mediated hydrogen bonding discussed above. Our substrate profiling results, which showed that the spSrtA_anthracis_ protein was inactive, are consistent with the idea that spSrtA and saSrtA enzymes have a different substrate-recognition mechanism than baSrtA (**Fig. S4B**).

#### (2) A non-covalent interaction between the β7-β8^−1^ residue and the β6^−2^ residue

We observe interactions between the β7-β8^−1^ and β6^−2^ residues in multiple Class A sortase structures (**Fig. 6B**). For example, K196 of saSrtA interacts with the β6^−2^ D160 in several of the states of the NMR structure, PDB code 2KID (**Fig. 6B**) (16). We also see a reasonable electrostatic interaction distance for the β7-β8^−1^ and β6^−2^ residues (K195 and E165, respectively) in several of the NMR states for *B. anthracis* SrtA (baSrtA) (PDB code 2RUI), as well as in the domain-swapped dimer structure of spSrtA, which shows the E214 β7-β8 loop residue of one protomer interacting with R184 (**Fig. 6B**). Interestingly, the nature of this interaction can change, e.g., in *Actinomyces oris* SrtA (aoSrtA) where both residues are hydrophobic leucine residues (L186 and L228 for the β6^−2^ and β7-β8^−1^ residues, respectively) (**Fig. 6B**).

Notably, our spSrtA homology model suggests that in spSrtA, the β7-β8^+1^ E208 may also interact with the β6^−2^ R184, with a distance between a guanidinium nitrogen atom of R184 and a side chain carboxylate oxygen atom on E208 equal to 2.7 Å (**Fig. S6A**). Because this residue is a glycine in the more active spSrtA_faecalis_ enzyme, we wanted to see if this β7-β8^+1^ Glu contributes to the relatively low reactivity of spSrtA by mutating the Gly in this position to Glu in our spSrtA_faecalis_ protein, or G145E (using *E. faecalis* SrtA numbering). We expressed and purified G145E spSrtA_faecalis_ and tested the protein with our A-, G-, and S-containing peptides. Indeed, we saw a 21-49% reduction in the activity for these three peptides relative to our initial spSrtA_faecalis_ variant, suggesting this residue may play a role in the activity of the enzyme (**Fig. 6C, Table S1**). The G145E spSrtA_faecalis_ protein is the only variant that contains less secondary structure content than the wild-type protein; therefore, we cannot rule out protein misfolding in these data (**Fig. S4A**). However, considering the β7-β8 loops of spSrtA and *E. faecalis* SrtA differ at only the β7-β8^+1^ (E in spSrtA, G in *E. faecalis* SrtA), β7-β8^+4^ (A versus Q, respectively), and β7-β8^−1^ (E versus T, respectively) positions (**Fig. 4A**), we hypothesize that this potential interaction at the β7-β8^+1^ position does negatively affect enzyme activity in the spSrtA and G145E spSrtA_faecalis_ proteins.

#### (3) A hydrophobic interaction between the β7-β8^+3^ residue and the β4-β5 loop (positions +2 or +3 from the catalytic His residue, or β4-β5^+2^/β4-β5^+3^)

In analyses of the previously published baSrtA structures, the authors of this work mention that the β4-β5^+2^/β4-β5^+3^ positions (for baSrtA, these are β4-β5^+2^ M128 and β4-β5^+3^ S129), play a role in stabilizing a hydrophobic residue, the β7-β8^+3^ V190, upon ligand binding (**Fig. 6D**) (33). Although we do not see a similar interaction in the bound saSrtA structure, we do see a similar movement of the β7-β8^+3^ residue towards these β4-β5 loop residues (**Fig. 6D**). The *L. monocytogenes* SrtA (or lmSrtA) structure (PDB ID 5HU4) also shows this interaction, β4-β5^+2^ M129 and β4-β5^+3^ R130 with P191 in the β7-β8^+3^ position, in the unbound state (**Fig. 6D**). Finally, our spSrtA model suggests a potential interaction between the β4-β5^+2^ I63 and β4-β5^+3^ F64 residues with β7-β8^+3^ L210 (**Fig. 6D**).

### Mutagenic investigation of the contribution of β7-β8 loop residues

In order to investigate these hypothesized interactions, we tested a number of β7-β8 mutants in spSrtA. Specifically, we expressed and purified the following mutant spSrtA proteins, as described in the Experimental Procedures, and characterized them using size exclusion chromatography and circular dichroism: E208G, L210P, T213A, D209T/T213D, and E214G (**Figs. 6E, S4A and S7C**). Our results are consistent with the proposed interactions. Specifically, the T213A mutation, which disrupts a potential intra-loop hydrogen bond between the β7-β8^+2^ D209 and β7-β8^+6^ T213 reduces spSrtA activity by 54-73% for G-, S-, and A-containing peptides (**Fig. 6E, Table S1**). When we attempt to reverse the hydrogen bond geometry with the D209T/T213D double mutant, we see no enzyme activity (**Fig. 6E, Table S1**). In contrast to an enhancement in activity when the intra-loop hydrogen bond is present, the β7-β8^−1^ and β6^−2^ proposed interaction negatively impacts spSrtA activity. The E214G mutant spSrtA revealed an almost 3-fold increase in activity, as compared to its wild-type counterpart (**Fig. 6E, Table S1**). This is consistent with recent work where a triple mutant of *S. pyogenes* SrtA (E189H/V206I/E215A, where E215A is a mutation at the equivalent β7-β8^−1^ position) resulted in a 6.6-fold enhanced catalytic efficiency (26). Notably, K196T in the catalytically enhanced pentamutant saSrtA protein is also located at the β7-β8^−1^ position (8). We also see that the β7-β8^+1^ Glu negatively affects enzyme activity, as we predicted based on our G145E spSrtA_faecalis_ results; here, the E208G spSrtA protein has over 2-fold higher activity than the wild-type protein (**Fig. 6E, Table S1**). We predict that in spSrtA, both E208 and E214 can interact with β6^−2^ R184, a hypothesis that our spSrtA model supports (**Fig. S6A**). Finally, the L210P spSrtA enzyme shows similar activity to the T213A mutant (**Fig. 6E, Table S1**), supporting a positive role for the proposed β7-β8^+3^ interaction with β4-β5^+2^/β4-β5^+3^, similar to that seen in baSrtA (33). This result may explain why our spSrtA_monocytogenes_ protein is catalytically inactive, as the *L. monocytogenes* SrtA β7-β8 loop contains a Pro at this position (**Figs. 4A, S4B**).

We tabulated the residues at relevant positions for the wild-type Class A sortases in the 8 organisms we used for our spSrtA chimeras in **Table 1**. Without structural data, we cannot conclude that all 3 interactions are present in all of these proteins, although most appear to be conserved at the sequence level. There are two notable exceptions, the sequences of *L. lactis* and *S. suis* SrtA proteins contain polar or charged residues at the β7-β8^−1^ position (H213 and Q213, respectively), but hydrophobic residues at the β6^−2^ residue (V183 and I183, respectively), which is inconsistent with a predicted interaction (**Table 1**). Homology models of both (template: 4TQX for *L. lactis* SrtA and 3FN5 for *S. suis* SrtA) suggest that the β6-β7^+1^ position in these proteins may be an interaction partner, although it is unclear (**Figs. S7D-E**). Although the other 8 SrtA sequences analyzed in **Table 1** do contain β6-β7^+1^ polar or charged residues (listed in the footnote of **Table 1**), no available SrtA structures show an interaction between this residue and the β7-β8 loop. Additional studies on these two proteins may elucidate whether or not this β7-β8 loop interaction is present and/or what, if any, effect there is on the activity and selectivity of the enzyme.

**Table 1.** The amino acid identities of the SrtA residues that are involved in β7-β8 loop-mediated interactions. Table 1 includes the amino acid positions in the wild-type proteins of all SrtA proteins discussed.

|  | *Interaction (1)* |  | *Interaction (2)* |  | *Interaction (3)* |  |  |
| --- | --- | --- | --- | --- | --- | --- | --- |
| | H-bond 1 | H-bond 2 | $\beta 6^{-2}$ | $\beta 7$-$\beta 8^{-1}$ | $\beta 4$-$\beta 5^{+2}$ | $\beta 4$-$\beta 5^{+3}$ | $\beta 7$-$\beta 8^{+3}$ |
| *B. anthracis* | S189<sup>a</sup> | D191<sup>a</sup> | E165 | K195 | M128 | S129 | V190 |
| *S. aureus* | N188 | T191 | D160 | K196 | F122 | I123 | Y187 |
| *E. faecalis* | D202 | T206 | K177 | T207 | T140 | E141 | L203 |
| *L. lactis* | D208 | T212 | V183<sup>b</sup> | H213 | M142 | T143 | A209 |
| *L. monocytogenes* | D189 | T194 | T166<sup>c</sup> | K196 | M129 | R130 | P191 |
| *S. oralis* | D213 | T217 | R188 | E218 | I147 | F148 | Y214 |
| *S. pneumoniae* | D209 | T213 | R184 | E214 | I143 | F144 | L210 |
| *S. suis* | D208 | T212 | I183<sup>d</sup> | Q213 | I142 | F143 | Y209 |

<sup>a</sup>*B. anthracis* SrtA: Although D191 appears to be positioned to interact with S189, the NMR structure PDB ID 2RUI (ordered loop in the baSrtA bound state) does not show hydrogen bonding between these residues in any of the 20 states.
<sup>b</sup>*L. lactis* SrtA: It appears more likely that H213 interacts with T185, the $\beta 6$ - $\beta 7^{+1}$ residue, based on a homology model (template: 4TQX, data not shown).
<sup>c</sup>*L. monocytogenes* SrtA: In the crystal structure (5HU4), T166 and K196 are not interacting.
<sup>d</sup>*S. suis* SrtA: It appears more likely that Q213 interacts with S185, $\beta 6$ - $\beta 7^{+1}$ residue, based on a homology model (template: 3FN5, data not shown). In the other proteins this position is: *B. anthracis* (T167), *S. aureus* (K162), *E. faecalis* (E179), *L. lactis* (T185), *L. monocytogenes* (D168), *S. oralis* (T190), and *S. pneumoniae* (T186).

### Sequence patterns in β7-β8 loop sequences of Class A sortases

Based on our structural observations using published structures and homology models, as well as our mutagenesis activity data, we identified 3 sequence motifs that appeared to be present in multiple Class A sortases. These include: (1) an intra-loop hydrogen-bonding pair, typically between the β7-β8^+1/+2^ loop and β7-β8^+6/+7^ positions, with the latter usually a Thr residue, (2) an interaction between the β7-β8^−1^ and β6^−2^ positions, and (3) an interaction between the β7-β8^+3^ with the β4-β5^+2^ and β4-β5^+3^ positions, usually of hydrophobic nature. In an effort to determine whether these motifs were more broadly present across the entire family of Class A sortases, we downloaded 387 Class A sortase sequences from the NCBI database. These were then aligned using MAFTT, and the β7-β8 loop sequences were manually extracted, using the catalytic Cys and Arg residues to define the boundaries of each (**Table S3**). Of those sequences, we analyzed a subset of 261 from the following genera: *Bacillus* (44 sequences), *Enterococcus* (68 sequences), *Lactobacillus* (102 sequences), *Listeria* (13 sequences), *Staphylococcus* (24 sequences), and *Streptococcus* (10 sequences).

We created WebLogos with sequences from each genus that had the same loop length and discovered a number of sequence patterns consistent with our predictions (**Figs. 7, S8**). Specifically, almost all of the sequences contain a β7-β8^+1/+2^ Asp/Glu paired with a β7-β8^+6/+7^ Thr residue, the potential stabilizing hydrogen bonding pattern described above (**Figs. 7, S8**). The two exceptions to this pattern are *Lactobacillus* sequences that are either 6 or 9 amino acids. In the 9 residue *Lactobacillus* sequences, however, there is a very strong prevalence of a β7-β8^+2^ Ser/Thr with a β7-β8^+7^ Glu, a residue pair which may also form a hydrogen bond. Consistent with this idea, in aoSrtA, there is an intra-loop hydrogen-bond between the β7-β8^+3^ S219 and β7-β8^+11^ D227 residues of *Actinomyces oris* SrtA (aoSrtA) (PDB ID 5UTT) (**Fig. S9A**).

**Fig. 7.**
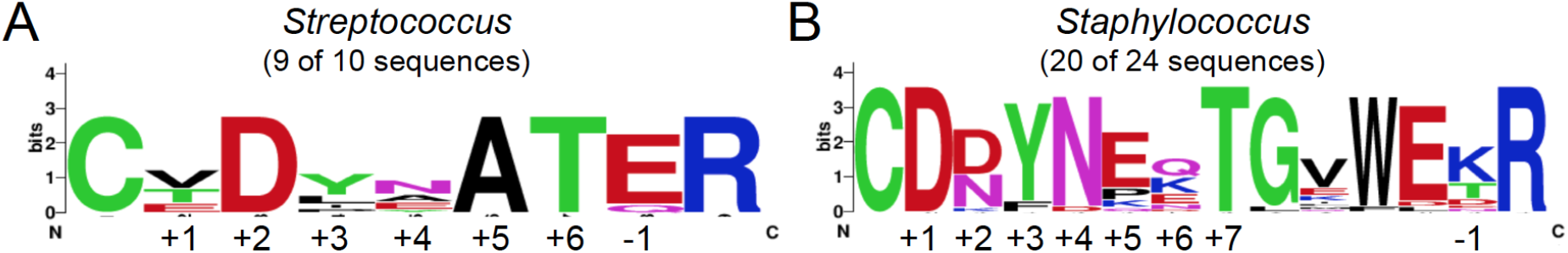
Conservation in the β7-β8 loop sequences of *Streptococcus* and *Staphylococcus* Class A sortase sequences. WebLogo analyses of β7-β8 loop sequences of the same lengths reveal conservation in the loops for both *Streptococcus* and *Staphylococcus* Class A sortases. The catalytic cysteine and arginine residues are also shown to provide reference points for the loop residues, which are labeled as in **Fig. 4A**. Loop sequences are in **Table S3**.

Second, of the 226 sequences that we included in our WebLogo analysis, only 32 do not contain a charged or polar residue in the β7-β8^−1^ position immediately preceding the catalytic Arg, including 19 *Lactobacillus* (Gly or Phe) and 13 *Bacillus* sequences (Gly or Trp) (**Table S3**). If we analyze these outliers further, we see that most often, including in *Lactobacillus pantheris*, which contains a Phe at this position, as well as in at least 8 *Lactobacillus* strains with a Gly at the β7-β8^−1^ position, we see Val, Phe, or Tyr residues at the β6^−2^ position, which is consistent with a favorable hydrophobic interaction between these residues (**Fig. S9B**). This interaction is seen in the aoSrtA structure, between residues β7-β8^−1^ L228 and β6^−2^ residue L186, described previously (**Fig. 6B**).

Finally, our WebLogo analysis confirms that the large majority of our sequences contain a β7-β8^+3^ hydrophobic (or proline) residue. In cases where this is not true (e.g., *Lactobacillus ozensis*, which has a β7-β8^+3^ Thr residue) the β4-β5^+2^ and/or β4-β5^+3^ residues are often also not strictly hydrophobic (e.g., for *L. ozensis*, the β4-β5^+3^ is a Glu) (**Fig. S9B**). Taken together, the three β7-β8 loop-mediated interactions described here appear to be broadly conserved in Class A sortases.

## Discussion

Although target sequence recognition by *S. aureus* SrtA is rigidly selective for a P1’ glycine, this is not true of all Class A sortases, such as those from *Streptococcus pneumoniae* and *Streptococcus pyogenes* (10, 30, 34, 35). Building from our previous work, in which spSrtA was found to accept peptides containing Gly-, Ala-, Ser- and other residues at P1’, we have shown here that this broadened substrate scope can be attributed to the sequence of the β7-β8 loop (26). Moreover, variations in β7-β8 loop sequences can substantially impact overall enzyme activity, affording chimeric sortases that outperform their wild-type counterpart i*n vitro*. Together with others, the present study implicates all of the variable loops in Class A sortases as being important determinants of enzyme function (13, 15, 18, 33).

With respect to structure, we propose three interactions that are facilitated by residues in the β7-β8 loop of spSrtA, characteristics which we suggest are broadly shared by Class A sortases. They are: (1) an intra-loop hydrogen bond that positively affects catalytic efficiency, typically mediated by a threonine residue at the β7-β8^+6^ or β7-β8^+7^ position, (2) an interaction that hinders enzyme activity between the β7-β8^−1^ and β6^−2^ residues, and (3) a positive interaction between the β7-β8^+3^ and β4-β5^+2^/β4-β5^+3^ residues, typically of hydrophobic nature. Notably, there appear to be other structural features in this structurally conserved loop that are unique to certain Class A sortases. These include the W194T residue of saSrtA, which specifically interacts with the P1 position of the CWSS (16). Others identified a disordered-to-ordered transition of the baSrtA β7-β8 loop, as well as regulation by an N-terminal appendage, although more research is needed to determine whether or not this is shared by other Class A sortases (33). Of the seven-residue β7-β8 loop in spSrtA, we did not specifically investigate the β7-β8^+4^ or β7-β8^+5^ positions, which are ^211^AA^212^ in spSrtA. Other *Streptococcus* SrtA structures, including those from *S. pyogenes* (^212^EA^213^), *S. agalactiae* (^188^EA^189^), and *S. mutans* (^209^GA^210^) show that these residues are interacting with solvent and often not well modeled in crystal structures (**Fig. S9C**). We reveal, however, that all other residues in the β7-β8 loop, e.g., the β7-β8^+1^ E208 residue in spSrtA, can affect activity, suggesting that it would be interesting to look at positional effects in Class A sortases with variable length β7-β8 loops.

In addition to informing our fundamental understanding of sortase substrate recognition, this work also has implications for the continued development of sortase-mediated ligation (SML) as a protein engineering tool (3, 36). Through exchange of β7-β8 loop residues between Class A sortases, we have generated chimeras such as spSrtA_faecalis_ and spSrtA_lactis_ with measurable activity against peptides possessing 15 of the 20 amino acids at P1’. With additional development, each of these sortase chimera/substrate combinations potentially offers a new handle for *in vitro* SML applications. While preliminary attempts here to modify a protein target displaying an LPAT**V** sequence using spSrtA_faecalis_ were unsuccessful, we consider it likely that optimization of the placement of the LPAT**V** site may restore reactivity. This includes examination of the accessibility requirements for the LPAT**V** sequence, and assessment of the impact of residues N- or C-terminal to the core LPAT**V** motif. Similar factors are known to affect the success of SML reactions with the widely used saSrtA/LPXT**G** system (37–39), and may need to be evaluated for our chimeras.

If successful, the development of these new sortase/substrate pairs has exciting consequences for SML engineering efforts: (1) it increases options for dual-labeling single proteins or multiplexed labeling of multiple proteins in the same systems (11, 40, 41), and (2) it may reduce the need to mutate naturally occurring protein sequences in order to render their termini compatible with SML. For example, using our previously published program, *MotifAnalyzer*, we found 190 instances of LPXT**G** in 189 unique proteins in the human proteome. However, if the P1’ position is now flexible, this number becomes 3606 instances of LPXT**X** in 2930 unique proteins (42).

Finally, the three variable loops mentioned here (β4-β5, β6-β7, and β7-β8) are conserved in all classes of sortases (**Fig. 1C**), and previous work determining and engineering sortase selectivity of different classes, e.g., sortase B, suggests similar roles for these loops in substrate recognition (5, 15, 43). Developing a deeper understanding of how residues in these loops affect substrate selectivity in all sortase classes may enable dramatic expansion of the sortase “toolbox” (**Fig. 8**), potentially allowing the development of ligases that are tailored to the needs of specific protein targets while also limiting off-target effects (5, 11, 13, 14, 27, 34, 44). In the over 20 years since saSrtA was discovered, the sortase superfamily has proven to be both a workhorse for protein engineering efforts, as well as an exciting system for future discoveries and insight into the stereochemistry and mechanisms of target recognition.

**Fig. 8.**
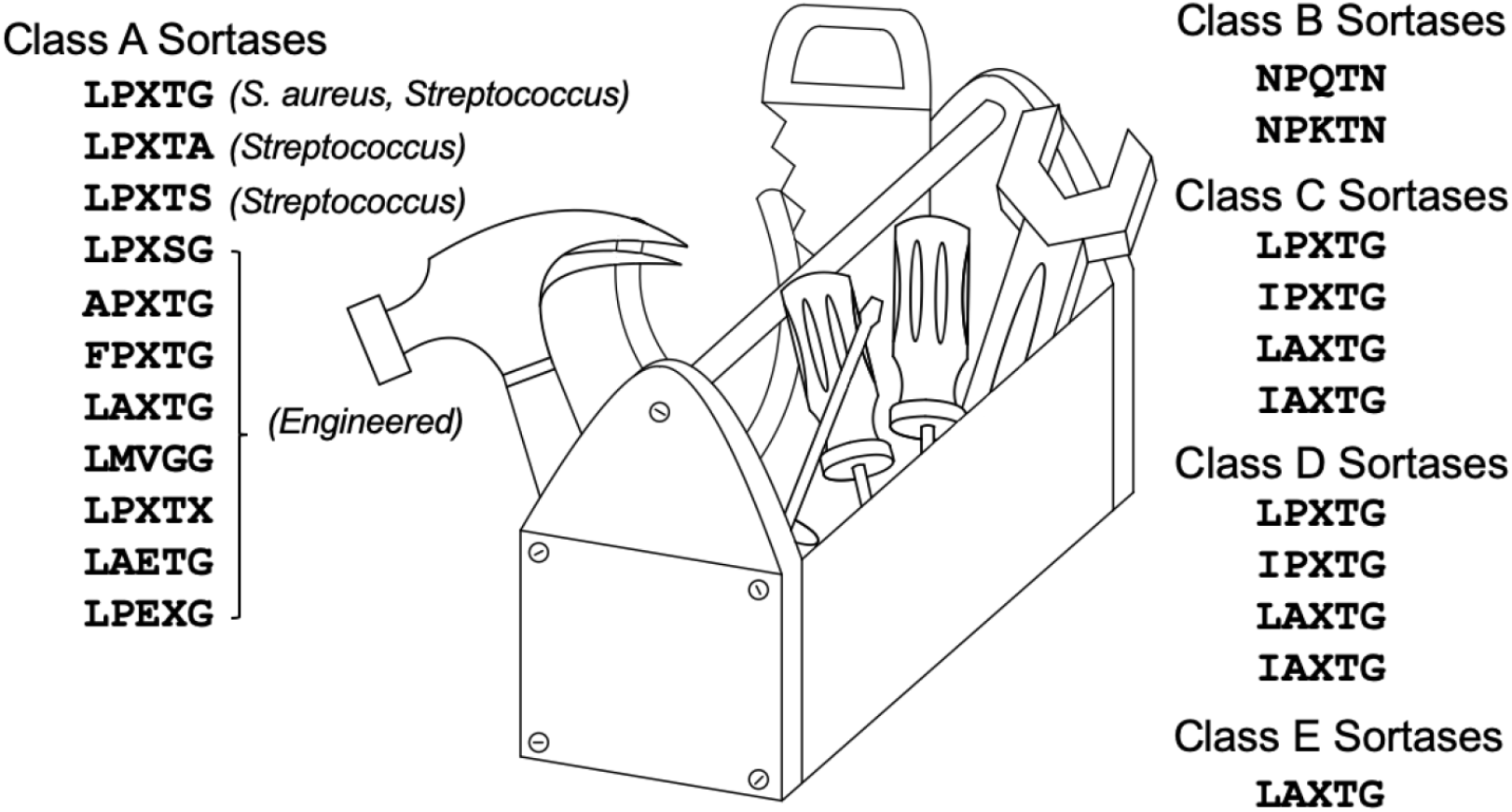
Building a sortase toolbox for SML experiments. Work from ourselves and others can be used to create a sortase “toolbox” for SML experiments, taking advantage of the various sequence motifs, both endogenous and engineered. Recognition sequences for various sortase subclasses are described in (5, 11, 13, 14, 27, 34, 44).

## Experimental Procedures

### Principal component analysis (PCA)

Initial sequences were obtained from UniProt and an alignment was generated by MAFFT (19, 20). Initially, each sequence was given a score for the number of gaps present for each residue and the filtered alignment was realigned by MAFFT. Subsequent analysis included all sequences without taking gaps into consideration (**Fig. 2B** versus **Figs. 2A, S2C**). The sortase multiple sequence alignment (MSA) was converted to a matrix representing the chemical properties of the amino acids, each amino acid was associated with 4 biochemical traits and a binary trait occupancy, as described. Each trait was normalized to the range from zero to one. In addition, gaps were given the average value of the matrix column with the exception of occupancy, so that they would not contribute to variance of the column. Gapped positions were given an occupancy score of zero (for the other chemical properties gapped positions received the average score). After translating the MSA, the resulting matrix was re-centered so that the matrix had a column-wise mean of zero. Principal component analysis was performed on the matrix by the singular value decomposition algorithm provided in the *scikit learn* Python package (45). Clustering was performed by a Gaussian mixture model provided in the *scikit-learn* Python package (45). Optimal cluster numbers were scored by Bayesian information criterion. Visualization was performed using a script written in Python with matplotlib.

### Protein expression and purification

Wild-type spSrtA and saSrtA proteins were expressed and purified as previously described (27). All other constructs, including chimeric and mutant proteins, were purchased from Genscript in the pET28a(+) vector. In general, protein expression and purification protocols were very similar to those previously described (27). Briefly, plasmids were transformed into *Escherichia coli* BL21 (DE3) competent cells and grown in LB media, with protein induction at OD_600_ 0.6-0.8 using 0.15 M IPTG for 18-20 h at 18°C.

Following cell harvest in lysis buffer [0.05 M Tris pH 7.5, 0.15 M NaCl, 0.0005 M ethylenediaminetetraacetic acid (EDTA)], the protein was purified using a 5 mL HisTrap HP column (GE Life Sciences, now Cytiva), using wash [0.05 M Tris pH 7.5, 0.15 M NaCl, 0.02 M Imidazole pH 7.5, 0.001 M TCEP] and elution [wash buffer, with 0.3 M Imidazole pH 7.5] buffers. Size exclusion chromatography (SEC) was conducted using a HiLoad 16/600 Superdex 75 column (GE Life Sciences, now Cytiva) in SEC running buffer [0.5 M Tris pH 7.5, 0.15 M NaCl, 0.001 M TCEP]. Purified protein corresponding to the monomeric peak was concentrated using an Amicon Ultra-15 Centrifugal Filter Unit (10,000 NWML) and analyzed by SDS-PAGE and analytical SEC (**Figs. S2, S7**). Protein not immediately used was flash frozen in SEC running buffer and stored at −80°C.

### Peptide synthesis

Detailed synthetic procedures are provided in the Supplemental Data. Briefly, all peptides were synthesized via manual Fmoc solid phase peptide synthesis (SPPS). Peptides were synthesized either individually or in tandem using Fmoc Rink amide MBHA resin or Synphase lantern solid supports. All other materials, including suitably protected Fmoc amino acids, and reagents for coupling, deprotection, and resin cleavage were obtained from commercial sources and used without further purification. All peptides were purified using RP-HPLC and their identities were confirmed via ESI-MS. Prior to use in sortase-catalyzed transacylation reactions, each purified peptide was prepared as a concentrated stock solution in DMSO and/or H_2_O (see Supplemental Data for details).

### Fluorescence Assay for Sortase Activity

Reactions were performed in a Costar round-bottom, black 96-well plate at a 100 μL reaction volume under the following conditions: 5 μM sortase, 50 μM peptide substrate, and 5 mM hydroxylamine nucleophile. All reactions contained 10% (*v/v*) 10x sortase reaction buffer (500 mM Tris pH 7.5, 1500 mM NaCl, and 100 mM CaCl_2_). Reactions also contained residual DMSO from the peptide stock solutions (0.5-1.5% (*v/v*), with the exception of the Phe- and Val-containing peptides at 5%). The peptides containing phenylalanine or valine required 5% (*v/v*) DMSO for solubility under the reaction conditions. 1 mM TCEP was also included in reactions utilizing the Abz-LPATCG-K(Dnp) substrate. Reactions were initiated by the addition of the sortase enzyme, which were prepared as 10x stock solutions in 50 mM Tris pH 7.5, 150 mM NaCl, and 1 mM TCEP. Microplates were analyzed using a Biotek Synergy H1 plate reader. The fluorescence intensity of each well was measured at 2-min time intervals over a 2-hr period at room temperature (λ_ex_ = 320 nm, λ_em_ = 420 nm, and detector gain = 75). All reactions were performed in triplicate (**Table S1**). For each substrate sequence, the background fluorescence of the intact peptide in the absence of enzyme was subtracted from the observed experimental data. Background-corrected fluorescence data was then normalized to the fluorescence intensity of a benchmark reaction between wild-type saSrtA and Abz-LPATGG-K(Dnp) (**Fig. S3A**).

### HPLC and LC-MS Characterization of Sortase-Catalyzed Reactions

Select pairings of sortase enzyme (5 μM, or 10 μM for the X-NH_2_ reactions), substrate (50 μM), and nucleophile (5 mM H_2_NOH or X-NH_2_) were repeated in the presence or absence of Ca^2+^ under reaction conditions that were otherwise identical to those described above for the fluorescence assay. These reactions were then analyzed using a Dionex Ultimate 3000 HPLC system interfaced with an Advion CMS expression^L^ mass spectrometer. Separations were achieved with a Phenomenex Kinetix^®^ 2.6 μM C18 100 Å column (100 x 2.1 mm) [aqueous (95% H_2_O, 5% MeCN, 0.1% formic acid) / MeCN (0.1% formic acid) mobile phase at 0.3 mL/min, method: hold 10% MeCN 0.0-0.5 min, linear gradient of 10-90% MeCN 0.5-7.0 min, hold 90% MeCN 7.0-8.0 min, linear gradient of 90-10% MeCN 8.0-8.1 min, re-equilibrate at 10% MeCN 8.1-13.25 min)].

### Circular dichroism spectroscopy

CD spectra were collected at 25°C on a Jasco J-1500 CD spectrometer. Samples were diluted to a calculated concentration of 0.2 mg/mL in 0.05 M Tris pH 7.5, 0.15 M NaCl, 0.001 M TCEP. After dilutions, concentrations were determined via absorbance at A280. Buffer subtracted measurements were taken in a 1 mm cuvette from 260 to 195 nm and represent an average of five scans. Each sample measurement was normalized for concentration and amino acid count.

### Sequence and structural analyses

All sequences were downloaded from either the NCBI database or UniProt, as indicated (19, 46, 47). Sequence alignments were performed using MAFFT, T-coffee, or BlastP (20, 48, 49). Alignments were visualized using Jalview (50). Homology modeling was performed using the SwissModel web interface (31, 32). Structural analyses and figure rendering were done using PyMOL.

## Acknowledgements

The authors would like to thank the other members of the Amacher and Antos labs for helpful discussions and assistance. JFA was funded by NSF CHE-1904711 and JMA was funded by a Cottrell Scholar Award from the Research Corporation for Science Advancement. KLH was supported by the National Institute of Allergy and Infectious Disease, Award Number F32AI145111. In addition, IMP received an Elwha Undergraduate Summer Research Award and DAJ received a Joseph & Karen Morse Student Research in Chemistry Fellowship to fund summer research. Start-up funds from Western Washington University also contributed to this project.

## Author contributions

IMP, SAS, JDV, KLH, JMA, and JFA designed the experiments. IMP, SAS, JDV, DAJ, MG, JES, HMK, and KLH performed experiments. KJ and EB contributed solid phase peptide synthesis to the project. IMP, SAS, JDV, DAJ, MG, KLH, JMA, and JFA contributed to data analysis. JMA and JFA wrote the manuscript. IMP, JDV, MG, KLH, JMA, and JFA prepared the figures. All authors contributed to editing of the manuscript.

