## Supplemental Material for "A second specificity-determining loop in Class A sortases: Biochemical characterization of natural sequence variation in chimeric SrtA enzymes"

### Table of Contents

**Fig S1.** Principal Component Analysis (PCA) of sortase superfamily 2

**Fig S2.** Representative analytical SEC chromatograms of sortase preparations following IMAC and preparative SEC, and circular dichroism (CD) spectra of wild-type sortase and variant proteins 3

**Table S2.** LC-MS characterization of synthetic peptides and reaction products 4

**Fig S3.** Sample benchmark reaction data and HPLC analysis of model reactions 5

**Fig S4.** Additional biochemical data for chimeric spSrtA proteins 6

**Fig S5.** Comparison of $spSrtA_{faecalis}$ substrate selectivity trends (HPLC vs fluorescence) 8

**Fig S6.** The homology model of spSrtA is very similar to other *Streptococcus* Class A structures 9

**Fig S7.** Structural characteristics of the β7-β8 loops of multiple Class A sortases 10

**Table S3.** Sequences of the β7-β8 loops of Class A sortases 12

**Fig S8.** WebLogo analyses of β7-β8 loop sequences in multiple genera 18

**Fig S9.** Sequence analysis of several SrtA proteins supports predicted β7-β8 interactions 19

Supplemental experimental procedures for peptide synthesis 20

References 24

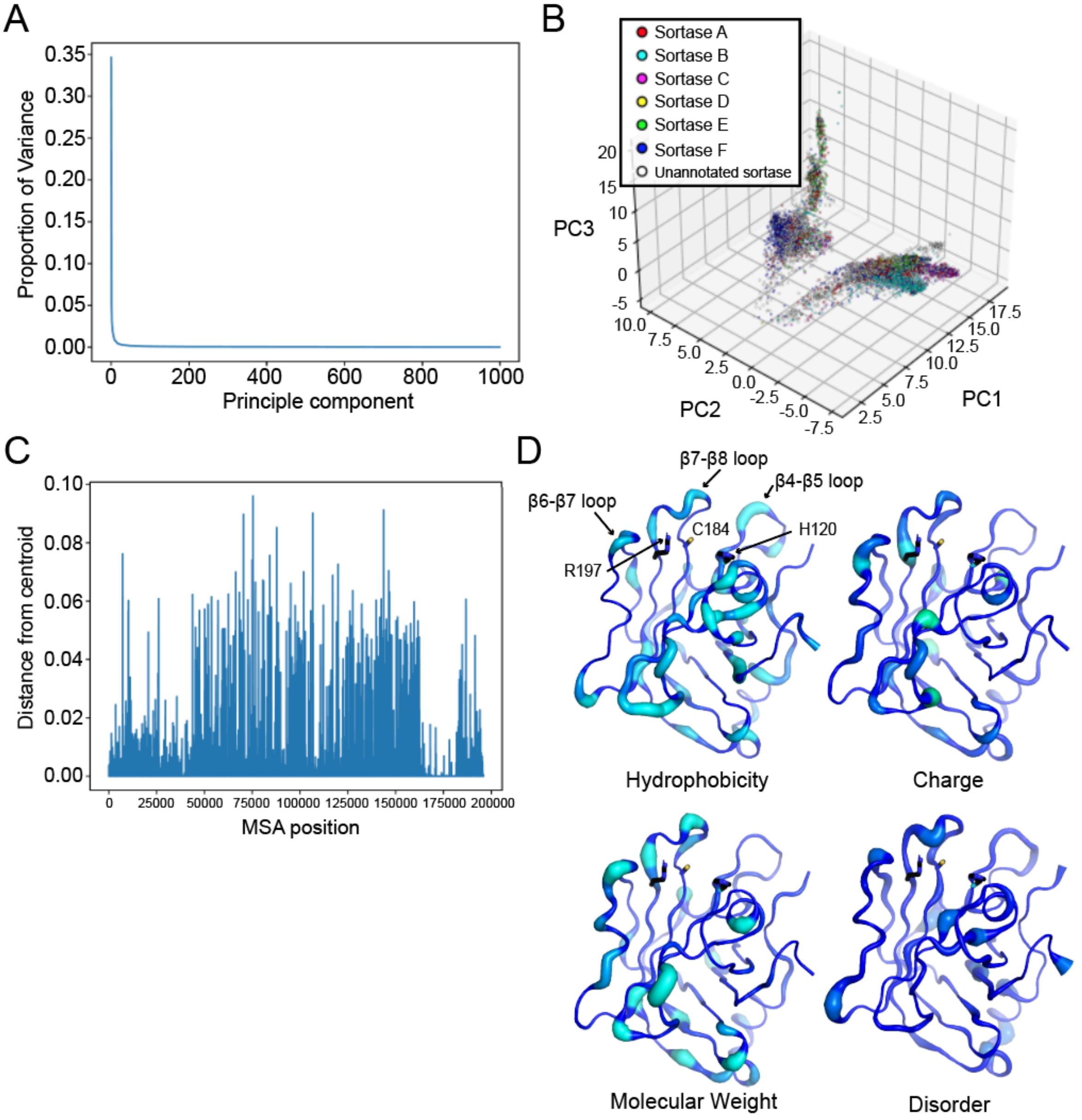


**Fig. S1. Principal component analysis (PCA) of sortase superfamily reveals sequence variability in structurally-conserved loops.** (A) Scree plot showing the variance explained for each principle component for the first 1000 principle components. (B) Scatter plot of all 39,188 sortase proteins available from UniProt in principle component space for the first three principal components PC1, PC2, and PC3. (C) Distance from the origin for each position in the multisequence alignment for the first three principle components. (D) Variable residues for hydrophobicity, charge, molecular weight and disorder propensity highlighted on PDB 3FN5, as labeled. The *S. pyogenes* SrtA protein is shown in cartoon representation. Both color and width indicate level of variability that resulted in the PCA, with lighter colors and greater width indicating a larger degree of variability.

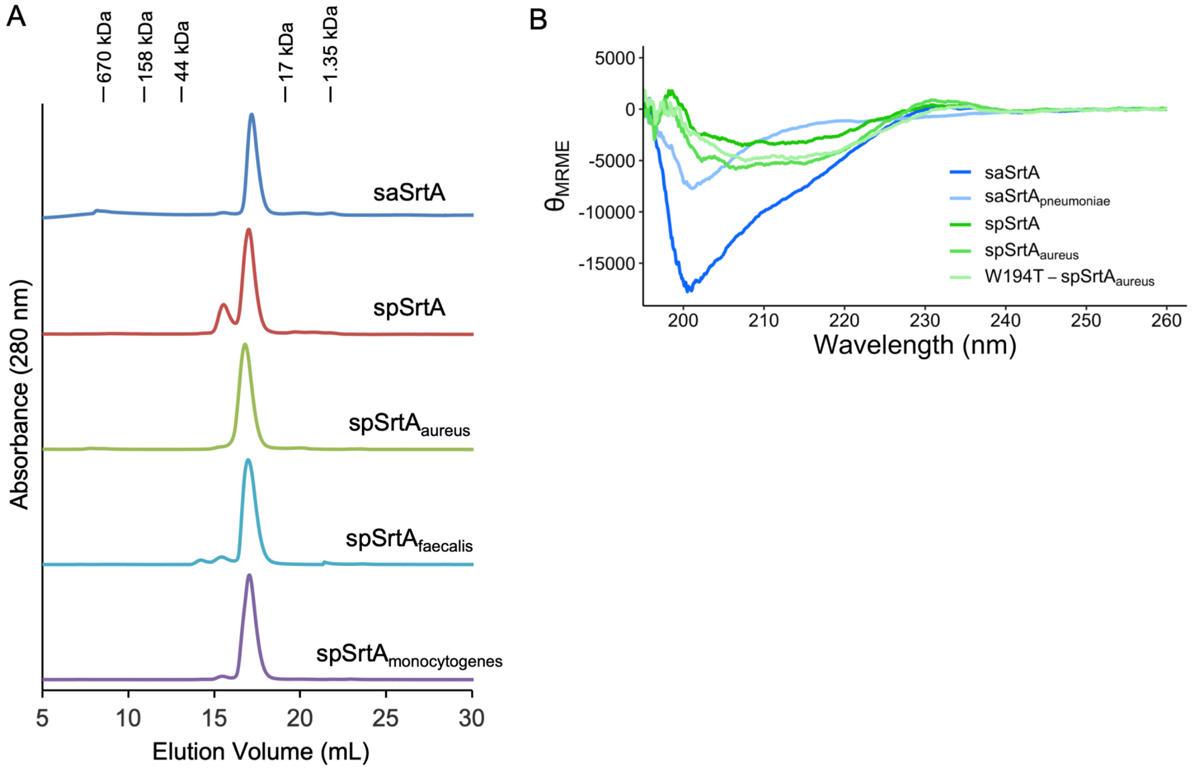


**Fig. S2. Representative analytical SEC chromatograms of sortase preparations following IMAC and preparative SEC, and circular dichroism (CD) spectra of wild-type sortase and variant proteins.** (A) Separations achieved using a Superdex 200 Increase 10/300 GL column (GE Life Sciences) with a mobile phase consisting of 0.5 M Tris pH 7.5, 0.15 M NaCl, 0.001 M TCEP. Elution volumes for molecular weight standards (Bio-Rad) are indicated above the chromatograms. (B) CD Spectra of saSrtA (blue) and spSrtA (green) indicating that the proteins vary in their secondary structure content. The chimeras, $saSrtA_{pneumoniae}$ (light blue), $spSrtA_{aureus}$ (mid-green), and W194T-$spSrtA_{aureus}$ (light green), retain much of the character of the parent protein, but gain some of the features of the protein from which the β7-β8 loops originate. Mean residue molar ellipticy ($\theta_{MRME}$) is in units of deg*cm$^2$/dmol.

**Table S2.** Mass spectrometry (LC-MS) characterization of synthetic peptides and relevant products from *in vitro* sortase-catalyzed model reactions.[a]

| Peptide | Mass (m/z) | |
|---|---|---|
| | **calculated** | **observed** |
| Abz-LPATAG-K(Dnp) | 941.4 | 941.5 |
| Abz-LPATCG-K(Dnp) | 973.4 | 973.3 |
| Abz-LPATDG-K(Dnp) | 985.4 | 985.5 |
| Abz-LPATEG-K(Dnp) | 999.5 | 999.4 |
| Abz-LPATFG-K(Dnp) | 1017.5 | 1017.5 |
| Abz-LPATGG-K(Dnp) | 927.4 | 927.5 |
| Abz-LPATHG-K(Dnp) | 1007.5 | 1007.4 |
| Abz-LPATIG-K(Dnp) | 983.5 | 983.5 |
| Abz-LPATKG-K(Dnp) | 998.5 | 998.6 |
| Abz-LPATLG-K(Dnp) | 983.5 | 983.6 |
| Abz-LPATMG-K(Dnp) | 1001.4 | 1001.4 |
| Abz-LPATNG-K(Dnp) | 984.5 | 984.5 |
| Abz-LPATPG-K(Dnp) | 967.5 | 967.6 |
| Abz-LPATQG-K(Dnp) | 998.5 | 998.4 |
| Abz-LPATRG-K(Dnp) | 1026.5 | 1026.5 |
| Abz-LPATSG-K(Dnp) | 957.4 | 957.3 |
| Abz-LPATTG-K(Dnp) | 971.5 | 971.5 |
| Abz-LPATVG-K(Dnp) | 969.5 | 969.4 |
| Abz-LPATWG-K(Dnp) | 1056.5 | 1056.6 |
| Abz-LPATYG-K(Dnp) | 1033.5 | 1033.4 |
| Abz-LPAT-NHOH | 535.3 | 535.3[b] |
| Abz-LPATA-$NH_2$ | 590.3 | 590.3[b] |
| Abz-LPATS-$NH_2$ | 606.3 | 606.3[b] |
| Abz-LPATS-$NH_2$ | 618.4 | 618.4[b] |
| AG-K(Dnp) | 440.2 | 440.2[b] |
| FG-K(Dnp) | 516.2 | 516.2 |
| GG-K(Dnp) | 426.2 | 426.2[b] |
| IG-K(Dnp) | 482.2 | 482.2 |
| LG-K(Dnp) | 482.2 | 482.2 |
| MG-K(Dnp) | 500.2 | 500.1 |
| NG-K(Dnp) | 483.2 | 483.2 |
| SG-K(Dnp) | 456.2 | 456.2[b] |
| VG-K(Dnp) | 468.2 | 468.2 |
| WG-K(Dnp) | 555.2 | 555.2 |
| YG-K(Dnp) | 532.2 | 532.2 |
| G-K(Dnp) | 369.2 | 369.2[b] |

[a]Calculated and observed masses represent $[M+H]^+$ ions (monoisotopic). [Abz = 2-aminobenzoyl fluorophore, Dnp = 2,4-dinitrophenyl chromophore, -NHOH = hydroxamic acid at C-terminus, -$NH_2$ primary amide at C-terminus]. [b]Product observed in multiple independent *in vitro* sortase-catalyzed model reactions. In all cases the observed m/z was within ±0.1 of the calculated m/z.

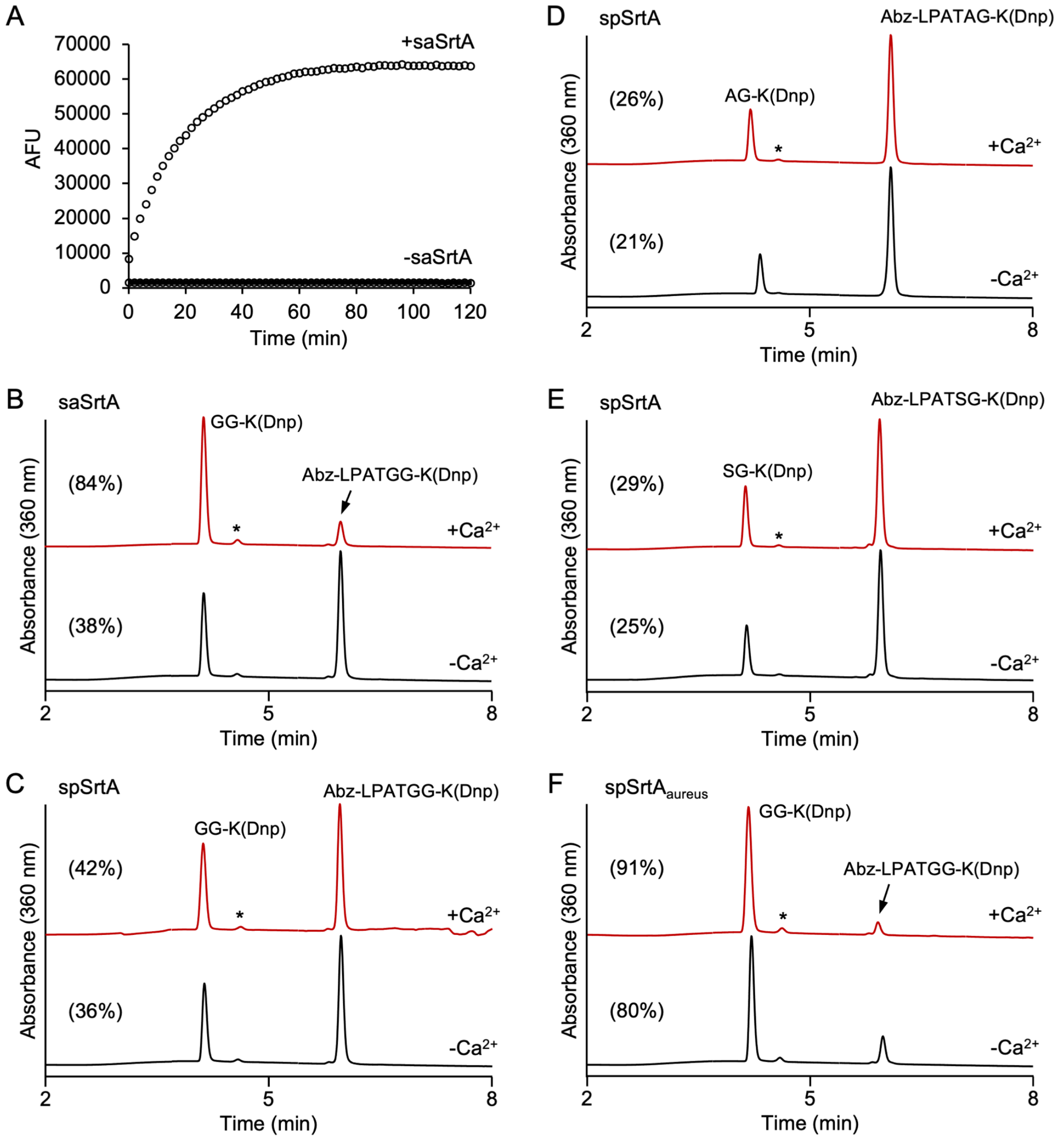


**Fig S3. Sample benchmark fluorescence data and HPLC characterization of select transacylation reactions in the presence and absence of $Ca^{2+}$.** (A) Representative fluorescence data ($\lambda_{ex}$ = 320 nm, $\lambda_{em}$ = 420 nm) for the reaction of Abz-LPATGG-K(Dnp) and $H_2NOH$ in the presence and absence of saSrtA. The benchmark AFU value used for scaling the majority of fluorescence data for other enzyme/substrate pairings was determined from six independent experiments. A benchmark AFU value derived from three additional, independent saSrtA/Abz-LPATGG-K(Dnp) reactions was used for scaling the fluorescence data in **Figure 6E**. (B-F) HPLC analyses of sortase-catalyzed reactions between Abz-LPATXG-K(Dnp) and $H_2NOH$ confirmed that the activity of saSrtA was $Ca^{2+}$-dependent (panel A), whereas spSrtA and $spSrtA_{aureus}$ did not require $Ca^{2+}$ (panels B-F). For all chromatograms, estimated substrate conversion at the 2 h timepoint is shown in parentheses. All peaks identities were confirmed via LC-MS (**Table S3**), and * denotes the position of the Abz-LPAT-NHOH ligation product. Low peak intensity is expected for this species due to the minimal absorbance of the Abz fluorophore at 360 nm.

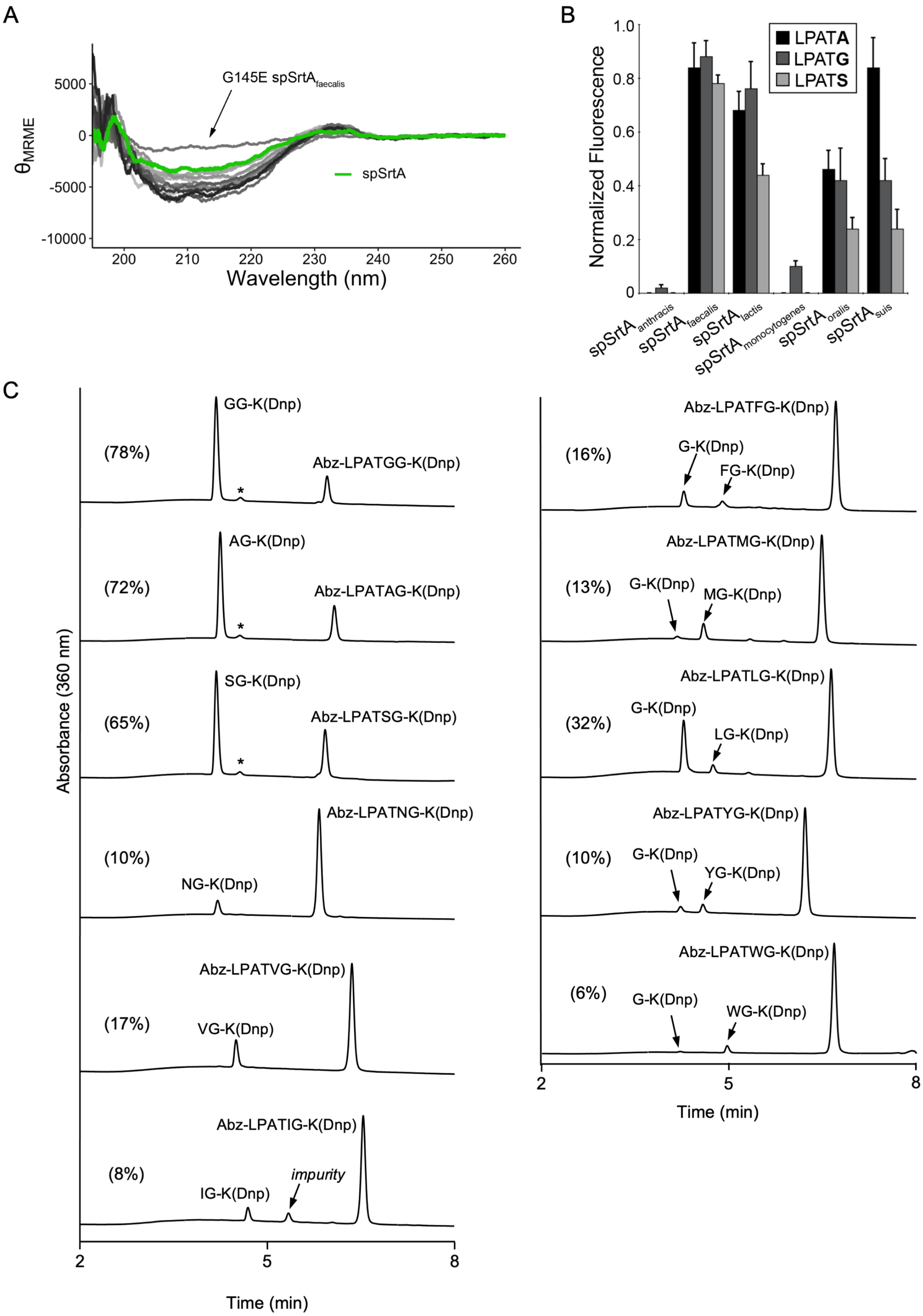

A
G145E spSrtA_faecalis
spSrtA
θMRME
5000
0
-5000
-10000
200
210
220
230
240
250
260
Wavelength (nm)
B
Normalized Fluorescence
1.0
0.8
0.6
0.4
0.2
0
LPATA
LPATG
LPATS
spSrtA_anthracis
spSrtA_faecalis
spSrtA_lactis
spSrtA_monocytogenes
spSrtA_oralis
spSrtA_suis
C
Absorbance (360 nm)
(78%)
GG-K(Dnp)
Abz-LPATGG-K(Dnp)
(72%)
AG-K(Dnp)
Abz-LPATAG-K(Dnp)
(65%)
SG-K(Dnp)
Abz-LPATSG-K(Dnp)
(10%)
NG-K(Dnp)
Abz-LPATNG-K(Dnp)
(17%)
VG-K(Dnp)
Abz-LPATVG-K(Dnp)
(8%)
IG-K(Dnp)
impurity
Abz-LPATIG-K(Dnp)
2
5
8
Time (min)
(16%)
G-K(Dnp)
FG-K(Dnp)
Abz-LPATFG-K(Dnp)
(13%)
G-K(Dnp)
MG-K(Dnp)
Abz-LPATMG-K(Dnp)
(32%)
G-K(Dnp)
LG-K(Dnp)
Abz-LPATLG-K(Dnp)
(10%)
G-K(Dnp)
YG-K(Dnp)
Abz-LPATYG-K(Dnp)
(6%)
G-K(Dnp)
WG-K(Dnp)
Abz-LPATWG-K(Dnp)
2
5
8
Time (min)

**Fig S4. Additional biochemical data for chimeric spSrtA proteins.** (A) With the exception of G145E-spSrtA$_{faecalis}$, all variants of spSrtA (overlaid gray/black spectra) used in this study contain secondary structure content comparable to the parent spSrtA (green) enzyme. (B) spSrtA chimeras with different β7-β8 loop sequences exhibit varied activity against a small panel of LPAT**X** substrates. (B) HPLC analyses of select reactions between Abz-LPATXG-K(Dnp), $H_2NOH$, and spSrtA$_{faecalis}$ in the absence of $Ca^{2+}$ reveal single cleavage sites for certain substrates (X = G, A, S, N, V, I) and a mixture of cleavage products for others (X = F, M, L, Y, W). Similar variations in cleavage sites have been observed previously for wild-type spSrtA (1). For all chromatograms, overall substrate conversion at the 2 h timepoint is shown in parentheses. All peaks identities were confirmed via LC-MS (**Table S3**), and where visible * denotes the position of the Abz-LPAT-NHOH ligation product.

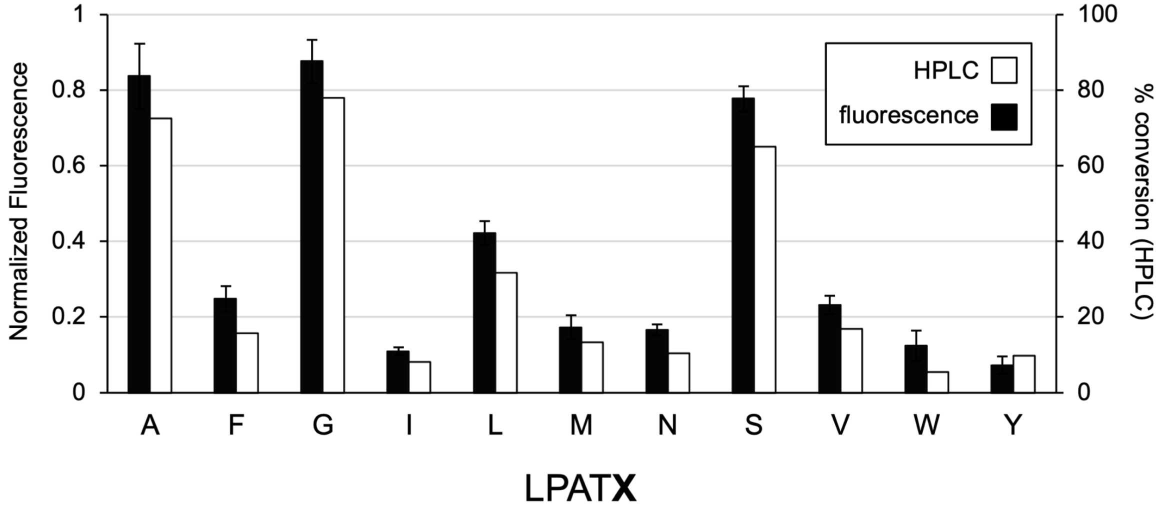


**Fig S5. Comparison of substrate selectivity trends for spSrtA$_{faecalis}$ as determined via HPLC and fluorescence assay.** Normalized fluorescence data (black) for the reaction of select Abz-LPATXG-K(Dnp) substrates with spSrtA$_{faecalis}$ is reproduced from **Fig. 4** in the main text, and was measured as described in Experimental Procedures. Percent substrate conversion, as determined by HPLC (white), was estimated from relevant peak areas observed in the 360 nm chromatogram. HPLC data represents single data points, while fluorescence experiments were conducted in triplicate. All data corresponds to the 2 h reaction timepoint.

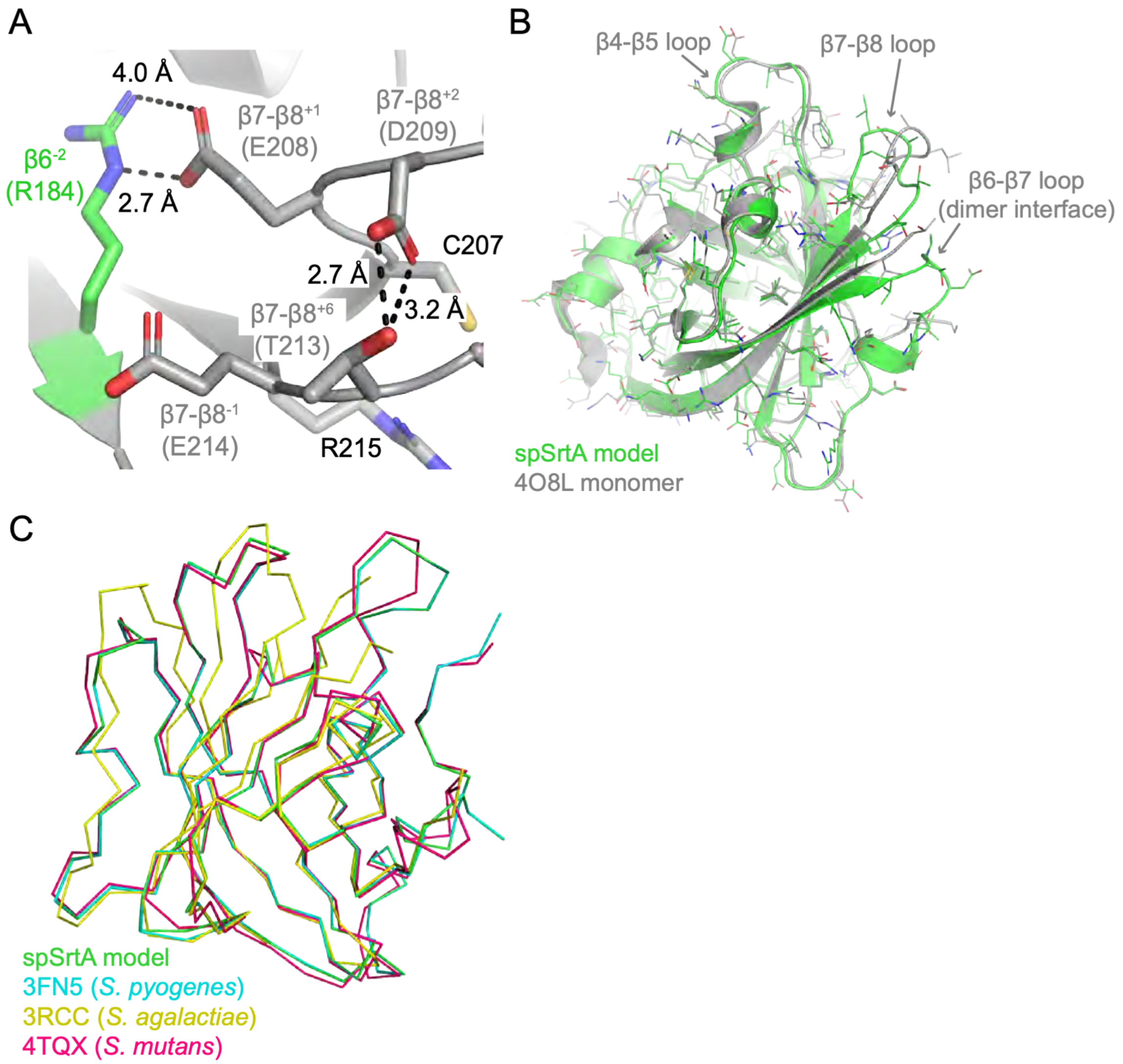


**Fig. S6. The homology model of spSrtA is very similar to other *Streptococcus* Class A structures.** (A) The spSrtA homology model was created using SwissModel and *Streptococcus pyogenes* SrtA (PDB ID 3FN5) as a template. This model (green cartoon), as well as (B) a monomer extracted from the domain-swapped dimer structure of spSrtA (4O8L, gray cartoon) are shown with the side chains as sticks and colored by heteroatom (O=red, N=blue, S=yellow). These two structures have an overall RMSD of 0.083 Å over 567 main chain atoms. Residues are perfectly aligned, with the exception of backbone variability in the β7-β8 loop and missing residues in the "4O8L monomer" β4-β5 loop, which is the location of the domain swapped region. Notably, the β7-β8 loop is also involved in the dimer interface in the 4O8L structure. (C) The ribbon traces of the spSrtA model (green), *S. pyogenes* SrtA (3FN5, cyan), *Streptococcus agalactiae* SrtA (3RCC, yellow), and *Streptococcus mutans* SrtA (4TQX, pink) are shown. Alignment of the model main chain atoms revealed overall RMSD values of 0.083 Å (567 atoms) for *S. pyogenes* SrtA, 0.773 Å (384 atoms) for *S. agalactiae* SrtA, and 0.456 Å (530 atoms) for *S. mutans* SrtA. Recall that the *S. pyogenes* SrtA structure was used as the model template, which explains the very low RMSD value.

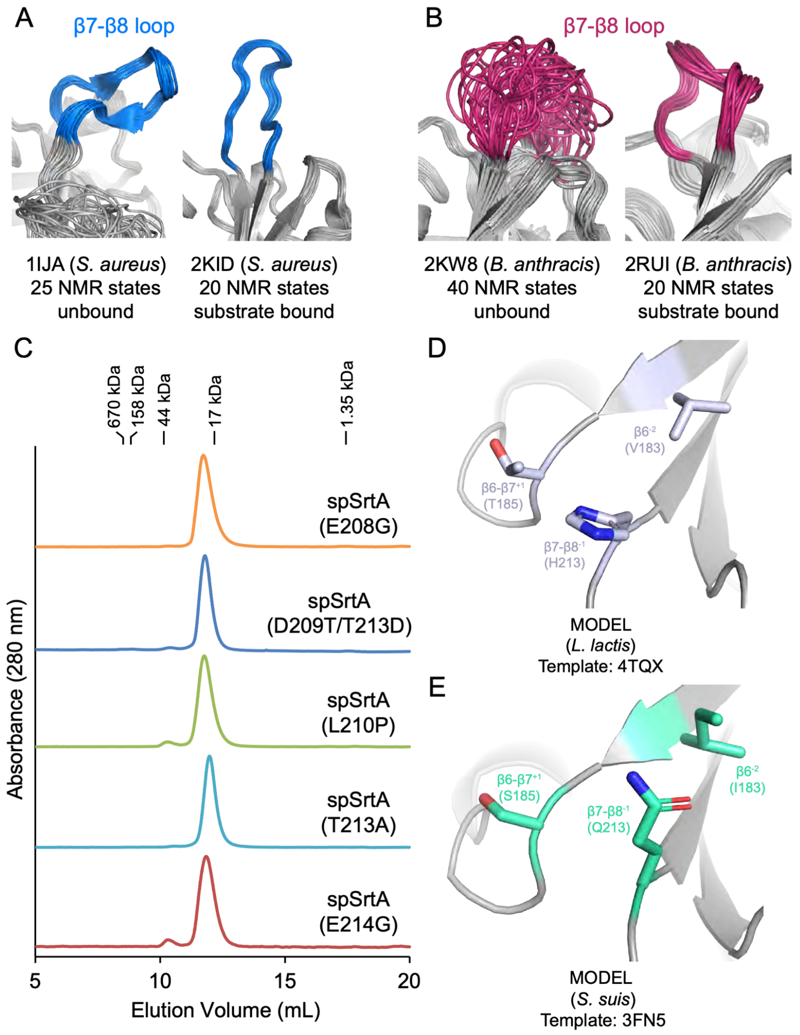


**Fig. S7. Structural characteristics of the β7-β8 loops of multiple Class A sortases.** (A-B) Flexibility in the β7-β8 loop differs in Class A sortases; for *S. aureus* SrtA (A), there is little to no flexibility in the loop in the unbound structure. In contrast, NMR structures of *B. anthracis* SrtA (B) reveal a large degree of flexibility in the unbound structure (left image, PDB ID 2KW8), which is reduced upon substrate binding (right image, 2RUI). For all, the proteins are in cartoon representation and colored gray, with the exception of the residues in the β7-β8 loop: *S. aureus* SrtA (blue) and *B. anthracis* SrtA (dark pink). (C) Separations achieved using an Enrich SEC 70 column (Bio-Rad) column with a mobile phase consisting of 0.5 M Tris pH 7.5, 0.15 M NaCl, 0.001 M TCEP at 0.5 mL/min. Elution volumes for molecular weight standards (Bio-Rad) are indicated above the chromatograms. (D-E) Homology models of *L. lactis* SrtA (A) and *S. suis* SrtA (B) are shown in gray cartoon representation, with residues in the β6 and β6-β7 loop that may interact with the β7-β8$^{-1}$ residue colored as labeled. Side chain sticks for these residues are in stick representation and colored by heteroatom (N=blue, O=red).

### Table S3. Sequences of the β7-β8 loops of Class A sortases.

| Organism (NCBI classification) | B7-B8 loop sequences |
|---|---|
| *Streptococcus agalactiae* | CTDPEATER |
| *Streptococcus mitis* | CEDLAATER |
| *Streptococcus sp. oral* | CVDYNATER |
| *Streptococcus sp. VT 162* | CVDYNATER |
| *Streptococcus sp. HMSC072D05* | CVDYNATER |
| *Streptococcus pneumoniae* | CSAATRTPNR |
| *Streptococcus_suis* | CTDYYATQR |
| *Streptococcus_oralis* | CVDYNATER |
| *Steptococcus pyogenes* | CTDIEATER |
| *Streptococcus_pneumoniae* | CEDLAATER |
| *Staphylococcus warneri* | CDNYNKNTGVWEKR |
| *Staphylococcus devriesei* | CDDYNENTGVWEKR |
| *Staphylococcus simiae* | CDNYNEKTGVWETR |
| *Staphylococcus haemolyticus* | CDDYNEQTGVWEKR |
| *Staphylococcus petrasii* | CDNYNEQTGIWEKR |
| *Staphylococcus equorum* | CDNFNEQTGVWENR |
| *Staphylococcus agnetis* | CDDYDEKTGKWLKR |
| *Staphylococcus simulans* | CDDYNPQTGEWETR |
| *Staphylococcus sp. HMSC13A10* | CDNYNKETGVWEKR |
| *Staphylococcus epidermidis* | CDDYNEETGVWETR |
| *Staphylococcus lugdunensis* | CDDYNEKTGVWEKR |
| *Staphylococcus aureus DAR1161* | CDDYNEKTGVWEKR |
| *Staphylococcus epidermidis* | CDDYNEETGKWETR |
| *Staphylococcus capitis* | CDDYNEKTGVWEKR |
| *Staphylococcus pettenkoferi* | CDKYNQQTGQWEKR |
| *Staphylococcus auricularis* | CDDFNPETGMWDTR |
| *Staphylococcus equorum* | CDNYNEQTGEWEDR |
| *Staphylococcus nepalensis* | CDNYNEQTGEWEDR |
| *Staphylococcus sciuri* | CDNYNPDTLLFEER |
| *Staphylococcus_aureus* | CDDYNEKTGVWEKR |
| *Staphylococcus epidermidis* | CQDLQARQR |
| *Staphylococcus sp. HMSC066C03* | CQDLQATQR |
| *Staphylococcus sciuri* | CSDLRATNR |
| *Staphylococcus sp. HMSC066G04* | CSDVKGTNR |
| *Listeria ivanovii* | CDKPTETTKR |
| *Listeria grayi* | CDKPTETDKR |
| *Listeria rocourtiae* | CDVATETNKR |
| *Listeria sp. 102* | CDKPTETTKR |
| *Listeria innocua* | CDKPTETTKR |
| *Listeria fleischmannii* | CDKPTETTKR |
| *Listeria floridensis* | CDKPTETSKR |
| *Listeria_monocytogenes* | CDKPTETTKR |
| *Listeria monocytogenes* | CDSSVDGTAGR |
| *Listeria sp. ILCC801* | CDKPTATTNR |
| *Listeria monocytogenes* | CSSERNTSKR |
| *Listeria grayi* | CDKGTATDYR |
| *Listeria aquatica* | CDKRTSTENR |
| *Enterococcus mundtii* | CDQVQQTSRR |
| *Enterococcus pallens* | CDTPRQTDQR |
| *Enterococcus rivorum* | CDKPSYTDRR |
| *Enterococcus dispar* | CDKPTQTQWR |
| *Enterococcus gilvus* | CDKPTQTKQR |
| *Enterococcus raffinosus* | CDKPTQTDQR |
| *Enterococcus sp. kppr6* | CDKPTRTDQR |
| *Enterococcus sp. HMSC064A12* | CDKPTHTEQR |
| *Enterococcus sp. 3H8_DIV0648* | CDKPTHTEQR |

| | |
|---|---|
| *Enterococcus sp. 6C8_DIV0013* | CDVSGANR |
| *Enterococcus aquimarinus* | CGEAAGVTR |
| *Enterococcus faecium EnGen0257* | CGDMDAVTR |
| *Enterococcus faecium* | CGDLAAVTR |
| *Enterococcus_faecalis* | CGDLQATTR |
| *Enterococcus sp. HMSC05C03* | CDSTNATSNR |
| *Enterococcus sp. 10A9_DIV0425* | CEGGLNTENR |
| *Enterococcus faecium* | CEGGLNTTQR |
| *Enterococcus faecium* | CEGGLNTDKR |
| *Enterococcus pallens ATCC BAA351* | CEGGLNTTKR |
| *Enterococcus sp. HMSC072H05* | CEGGLNTPSR |
| *Enterococcus sp. HMSC05C03* | CEGAINTPNR |
| *Enterococcus avium* | CEGGLNTPKR |
| *Enterococcus gilvus* | CEGGLYTANR |
| *Enterococcus gilvus* | CEGGINTPNR |
| *Enterococcus faecium* | CEGGLHTPNR |
| *Enterococcus faecium* | CEGGLHTANR |
| *Enterococcus avium* | CDSSNHNTPNR |
| *Enterococcus malodoratus ATCC 43197* | CDSSNQNTPNR |
| *Enterococcus sp. 4E1_DIV0656* | CDGSRVGTDYR |
| *Enterococcus mundtii* | CDGSRAGTDYR |
| *Enterococcus faecium* | CDRPAVHTPNR |
| *Enterococcus faecalis* | CDHAVPGTNNR |
| *Enterococcus faecium* | CDSSVAGTNGR |
| *Enterococcus faecium* | CDSSIDGTDGR |
| *Enterococcus faecium LA4B2* | CDSSVAGTEGR |
| *Enterococcus villorum* | CDSSVAGTDGR |
| *Enterococcus mundtii* | CDKYEETNKR |
| *Enterococcus sulfureus* | CPTPVVTSQR |
| *Enterococcus sp. 8G7_MSG3316* | CPNARRSPHR |
| *Enterococcus columbae* | CEGGINTDNR |
| *Enterococcus italicus DSM 15952* | CVPDGKEVPDKR |
| *Enterococcus faecalis EnGen0327* | CYDDSTKLPENR |
| *Enterococcus faecalis* | CYDDNTKLPENR |
| *Enterococcus faecium EnGen0305* | CYDDNTKLPENR |
| *Enterococcus malodoratus* | CDSPDYTTKR |
| *Enterococcus pallens ATCC BAA351* | CDKPTLTKKR |
| *Enterococcus sp. 3H8_DIV0648* | CDRGTQTTGR |
| *Enterococcus faecium* | CDSVQATDQR |
| *Enterococcus mundtii* | CQTVQDTPNR |
| *Enterococcus hirae* | CQSVQTTDNR |
| *Enterococcus mundtii* | CQTVQTTDNR |
| *Enterococcus casseliflavus* | CPTPTRTEER |
| *Enterococcus gallinarum* | CPTPSRTDER |
| *Enterococcus phoeniculicola* | CPVPQSTKQR |
| *Enterococcus sp. 3H8_DIV0648* | CDEPTITDQR |
| *Enterococcus mundtii* | CDQPTLTDKR |
| *Enterococcus thailandicus* | CDKPQRTDKR |
| *Enterococcus faecalis EnGen0332* | CDQETETTGR |
| *Enterococcus faecalis* | CDQATKTTGR |
| *Enterococcus faecalis 02MBP10* | CDQATKTTGR |
| *Enterococcus faecalis RP2S4* | CDQATKTTGR |
| *Enterococcus faecalis* | CDKATKTTGR |
| *Enterococcus faecalis ATCC 6055* | CDQATKTDGR |
| *Enterococcus faecalis NY9* | CDQATKTDGR |
| *Enterococcus faecium* | CEGGTGTNYR |
| *Enterococcus massiliensis* | CEGGIGTEYR |
| *Enterococcus sp. 6C8_DIV0013* | CEGDIGTIYR |
| *Enterococcus cecorum* | CQEDAQFWNTYYRTGRTYAFYR |

| | |
|---|---|
| *Bacillus toyonensis* | CFGGLNTDKR |
| *Bacillus thuringiensis IBL 4222* | CFGGLNTDKR |
| *Bacillus cereus MC118* | CDKATLTDRR |
| *Bacillus cereus* | CDTPTLTDQR |
| *Bacillus anthracis* | CDKATTTNQR |
| *Bacillus cereus* | CDKATATNHR |
| *Bacillus manliponensis* | CITIKNNAKR |
| *Bacillus sp. AFS018417* | CSSVLDNSKR |
| *Bacillus pseudomycoides* | CVSVSDNSKR |
| *Bacillus cereus* | CVSVKDNSKR |
| *Bacillus_anthracis* | CVSVKDNSKR |
| *Bacillus megaterium* | CDIPSKPESR |
| *Bacillus pseudomycoides* | CLSIKDNSKR |
| *Bacillus cereus* | CLSIKDNSKR |
| *Bacillus toyonensis* | CYDDAGTTR |
| *Bacillus sp. 491mf* | CYDDKGETR |
| *Bacillus atrophaeus* | CDKAVETEGR |
| *Bacillus subtilis* | CDKAVKTEGR |
| *Bacillus subtilis* | CDKPTATEKR |
| *Bacillus sp. JCM 19035* | CDISGPTDQR |
| *Bacillus sp. JCM 19041* | CDISKPTNMR |
| *Bacillus sp. MarseilleP3800* | CDTSQPTTNR |
| *Bacillus simplex* | CDVTGIDTDKR |
| *Bacillus haynesii* | CNVSGIKTDKR |
| *Bacillus cereus* | CDISGPTNKR |
| *Bacillus cereus TIAC219* | CDVAGATDKR |
| *Bacillus cereus VD045* | CDVAEATDKR |
| *Bacillus pumilus* | CDLPTATTHR |
| *Bacillaceae bacterium* | CDVPTKTNKR |
| *Bacillus amyloliquefaciens* | CDKAVRTEGR |
| *Bacillus subtilis* | CDKAVKTEGR |
| *Bacillus atrophaeus* | CDKAVETEGR |
| *Bacillus sp. BSC154* | CDKAVETEGR |
| *Bacillus paralicheniformis* | CDKAVRTEGR |
| *Bacillus glycinifermentans* | CDKPERTKGR |
| *Bacillus sp. NSP9.1* | CDKAERTDRR |
| *Bacillus swezeyi* | CDKAERTEGR |
| *Bacillus lonarensis* | CYSEDGSDR |
| *Bacillus cereus* | CTNNGKKR |
| *Bacillus cereus* | CNANGKKR |
| *Bacillus cereus* | CNTNGKKR |
| *Bacillus cereus* | CNADGKKR |
| *Bacillus cereus* | CHAKGEDR |
| *Bacillus cecembensis* | CAEDGTRR |
| *Lactobacillus secaliphilus* | CDATGAR |
| *Lactobacillus uvarum* | CLTAKTGENNR |
| *Lactobacillus saerimneri* | CASGEPGETNR |
| *Lactobacillus johnsonii* | CTNDNKKR |
| *Lactobacillus frumenti* | CTSDNKRR |
| *Lactobacillus oris* | CTSDNQKR |
| *Lactobacillus sakei* | CWSPNHENNPKHR |
| *Lactobacillus sakei* | CDPDGEIVNGQTYER |
| *Lactobacillus iners* | CDYTGAGR |
| *Lactobacillus crispatus* | CDWTGQGR |
| *Lactobacillus bombicola* | CDYTGQGR |
| *Lactobacillus kimbladii* | CDWTGQGR |
| *Sporolactobacillus vineae* | CDKPTRTPNR |
| *Sporolactobacillus terrae* | CDKPTRTPNR |
| *Sporolactobacillus laevolacticus* | CDKPTRTPKR |

| | |
|---|---|
| *Sporolactobacillus nakayamae* | CDKPTRTPNR |
| *Lactobacillus fermentum* | CDYTGSHR |
| *Lactobacillus reuteri* | CDATGANR |
| *Lactobacillus rossiae* | CDATGANR |
| *Lactobacillus ozensis* | CDATGVGR |
| *Lactobacillus senioris* | CDATGAGR |
| *Lactobacillus farraginis* | CDATGKGR |
| *Lactobacillus buchneri* | CDATGKGR |
| *Lactobacillus kisonensis* | CDATGAGR |
| *Lactobacillus parakefiri* | CDATGAGR |
| *Lactobacillus parabuchneri* | CDATGAGR |
| *Lactobacillus paucivorans* | CDATGANR |
| *Lactobacillus zymae* | CDATGANR |
| *Lactobacillus kimchicus* | CDATGAGR |
| *Lactobacillus collinoides* | CDATGAGR |
| *Lactobacillus odoratitofui* | CDATGARR |
| *Lactobacillus malefermentans* | CDATGARR |
| *Lactobacillus oligofermentans* | CDATGEGR |
| *Lactobacillus vaccinostercus* | CDAHGKNR |
| *Lactobacillus hokkaidonensis* | CDATGTNR |
| *Lactobacillus wasatchensis* | CDATGTNR |
| *Lactobacillus fermentum* | CDATGARR |
| *Lactobacillus ingluviei* | CDATGANR |
| *Lactobacillus equigenerosi* | CNSDGSRR |
| *Lactobacillus secaliphilus* | CDATGARR |
| *Lactobacillus oris* | CNANGERR |
| *Lactobacillus sp. MarseilleP3519* | CDATGANR |
| *Lactobacillus frumenti* | CDATGANR |
| *Lactobacillus plantarum* | CDKTGAGR |
| *Lactobacillus rhamnosus* | CTADSQHR |
| *Lactobacillus aviarius* | CDQTNQKR |
| *Lactobacillus fuchuensis* | CEGALNTPNR |
| *Lactobacillus brevis* | CYEIPPDYANAQNR |
| *Lactobacillus ruminis* | CASGRVNEKRR |
| *Lactobacillus saniviri* | CASSQLDEANR |
| *Lactobacillus paucivorans* | CATAKRGEQNR |
| *Lactobacillus senmaizukei* | CASAKRNEPNR |
| *Lactobacillus koreensis* | CNSARRGEPKR |
| *Lactobacillus hammesii* | CASAKRGEPKR |
| *Lactobacillus parabrevis* | CNSAKRGEPKR |
| *Lactobacillus ruminis* | CASGMTGESRR |
| *Lactobacillus hordei* | CDPIKGVAHTPLR |
| *Lactobacillus vini* | CDPIKGVAHTPLR |
| *Lactobacillus ghanensis* | CDPVPGVARTPLR |
| *Lactobacillus nagelii* | CDPIPGVARTPLR |
| *Lactobacillus aquaticus* | CLTAKTGENNR |
| *Lactobacillus satsumensis* | CLTATTGETNR |
| *Lactobacillus oeni* | CLTATAGETNR |
| *Lactobacillus floricola* | CADGGVNR |
| *Lactobacillus iners* | CANGGKMR |
| *Lactobacillus amylophilus* | CANGGISR |
| *Lactobacillus iners* | CADWGANR |
| *Lactobacillus delbrueckii* | CADGGVNR |
| *Lactobacillus equicursoris 66c* | CADGGTNR |
| *Lactobacillus kefiranofaciens* | CADGGANR |
| *Lactobacillus crispatus* | CADGGKNR |
| *Lactobacillus pasteurii* | CADGGVNR |
| *Lactobacillus hamsteri* | CADGGVNR |
| *Lactobacillus jensenii JVV16* | CADGGINR |

| | |
|---|---|
| *Lactobacillus gasseri* | CADGGVNR |
| *Lactobacillus hominis* | CADGGVNR |
| *Lactobacillus senioris* | CASWRWNEPDR |
| *Lactobacillus mindensis* | CADGGTNR |
| *Lactobacillus ginsenosidimutans* | CADGGTNR |
| *Lactobacillus camelliae* | CEVSTASRADR |
| *Lactobacillus pantheris* | CASSLAGEEDR |
| *Lactobacillus selangorensis* | CSSATEGETNR |
| *Lactobacillus paracasei* | CDSATPNTPKR |
| *Lactobacillus saniviri* | CASPTEGEVNR |
| *Lactobacillus brantae* | CASPTEGETDR |
| *Lactobacillus casei* | CASPTEGEVDR |
| *Lactobacillus homohiochii* | CASGNPGETRR |
| *Lactobacillus kunkeei* | CASGTPDEPNR |
| *Lactobacillus parabuchneri* | CASAKTGEKNR |
| *Lactobacillus parabuchneri* | CASGKPEESRR |
| *Lactobacillus hokkaidonensis JCM 18461* | CDRAYGTDSR |
| *Lactobacillus wasatchensis* | CDRSYGTDSR |
| *Lactobacillus kefiri* | CASGKPEESNR |
| *Lactobacillus farraginis* | CASGKPEESNR |
| *Lactobacillus diolivorans* | CLTASIGESKR |
| *Lactobacillus harbinensis* | CEGPRGTDYR |
| *Lactobacillus plantarum* | CNATGSMR |
| *Lactobacillus shenzhenensis* | CSGGYDTPYR |
| *Lactobacillus brantae* | CDQPTATTGR |
| *Lactobacillus nasuensis* | CEGDVGTNFR |
| *Lactobacillus thailandensis* | CEGPLNTPFR |
| *Lactobacillus pantheris* | CEGPLNTPFR |
| *Lactobacillus brantae* | CEGPEGTPYR |
| *Weissella oryzae* | CNYTADNGR |
| *Weissella viridescens* | CDYTAERGR |
| *Weissella ceti* | CNYTAEAGR |
| *Weissella halotolerans* | CDYTAERGR |
| *Weissella confusa* | CDYTAERGR |
| *Weissella oryzae* | CFEEYPDYYHAKYR |
| *Weissella jogaejeotgali* | CLFPSTSYR |
| *Weissella confusa* | SLFPSTQYR |
| *Weissella kandleri* | CLFPSTEYR |
| *Weissella paramesenteroides* | CLFPSTDYR |
| *Weissella cibaria* | CDEPGLFTLHPENR |
| *Weissella oryzae SG25* | CDEEERWDTNTKSR |
| *Weissella kandleri* | CDERDEQKFNLSPVNR |
| *Mycobacterium abscessus* | CDNYNQQTGVWEKRK |
| *Bacillaceae bacterium* | *CDVPTKTSKRV* |
| *Brevibacterium halotolerans* | *CDKAVETEGR* |
| *Gemella morbillorum* | *CDNYNPKTGEWESR* |
| *Gemella asaccharolytica* | *CDGYNSVTGEWEER* |
| *Auricoccus indicus* | *CDDYDPNTGLFLTR* |
| *Halolactibacillus halophilus* | *CEGEYNTDWR* |
| *Aerococcus viridans* | *CEGGYGTDYR* |
| *Leuconostoc lactis* | *CSSAQRTPNR* |
| *Lactococcus lactis* | *CNQTLKTPYR* |
| *Lactococcus garvieae* | *CNTATQTPYR* |
| *Lactococcus garvieae* | *CNSATETPYR* |
| *Carnobacterium divergens* | *CSTDAGVER* |
| *Carnobacterium divergens* | *CSTDAGVER* |
| *Carnobacterium pleistocenium* | *CTQAGSKR* |
| *Aerococcus christensenii* | *CSKPVQPVHTR* |
| *Aerococcus urinae* | *CEESTSAAMR* |

| | |
|---|---|
| *Aerococcus christensenii* | CGESTSTAQR |
| *Aerococcus sp. HMSC10H05* | CTADLTQR |
| *Aerococcus sp. 1KP2016* | CTADLTQR |
| *Rhodococcus sp. 15238811a* | CDRATSTEYR |
| *Carnobacterium maltaromaticum* | CDVSSATNQR |
| *Carnobacterium divergens* | CDVPFQTDQR |
| *Carnobacterium divergens* | CDVPFQTDKR |
| *Brachybacterium faecium* | CDTADATSKR |
| *Brochothrix campestris* | CDTASATTKR |
| *Domibacillus antri* | CYKSSEPEKR |
| *Domibacillus enclensis* | CFDSSDREKR |
| *Domibacillus robiginosus* | CYSSDDRTKR |
| *Allofustis seminis* | CVYEKNKGFGFSDTGNRR |
| *Carnobacterium sp.* | CYYIDGQNSGDR |
| *Atopostipes suicloacalis* | CYYVDGKNSGDR |
| *Aerococcus suis* | CFYPQRYFDGDDDR |
| *Anaerosphaera aminiphila* | CYYENGKNTGNR |
| *Catonella morbi* | CDKGTWTSNR |
| *Varibaculum timonense* | CYYTSKNGKR |
| *Tissierella sp.* | CYYSSKTGKR |
| *Anaerosalibacter massiliensis* | CYHSSKTGKR |
| *Clostridium ultunense* | CYFSSKTGKR |
| *Criibacterium bergeronii* | CYFSSSTGKR |
| *Peptoanaerobacter stomatis* | CYYSSKTGKR |
| *Eubacterium yurii* | CYYSSNTGKR |
| *Proteiniclasticum ruminis* | CYYSSSTGKR |
| *Clostridium amylolyticum* | CYYSSKTGKR |
| *Peptostreptococcus sp.* | CNNDGSKR |
| *Amphibacillus sediminis* | CYSHDGSDR |
| *Alkalibacterium sp. AK22* | CNHDGSER |
| *Alkalibacterium gilvum* | CNHDGTER |
| *Marinilactibacillus psychrotolerans* | CNHDGTER |
| *Carnobacterium sp. AT7* | CNYDGTER |
| *Kurthia senegalensis* | CNFQGKKR |
| *Kurthia sp. 11kri321* | CNYDGSKR |
| *Viridibacillus arvi* | CNYDGSKR |
| *Rummeliibacillus stabekisii* | CNADGTQR |
| *Carnobacterium sp. CP1* | CNLTEQQR |
| *Pediococcus damnosus* | CDATGANR |
| *Pediococcus claussenii* | CDATGAKR |
| *Pediococcus acidilactici* | CDATGAGR |
| *Pediococcus pentosaceus* | CDDTGAGR |
| *Granulicatella sp. HMSC31F03* | CDDYNATKR |
| *Alkalibacterium gilvum* | CNNEGETR |
| *Globicatella sp. HMSC072A10* | CDYDLVER |
| *Dolosicoccus paucivorans* | CDTGLVDR |
| *Facklamia sourekii* | CDYGLVDR |
| *Bavariicoccus seileri* | CNVTGSKR |
| *Carnobacterium maltaromaticum* | CNQDGSKR |
| *Paenibacillus thiaminolyticus* | CDDSGKAR |
| *Lactococcus_lactis* | CADAEATHR |
| *Acetinomyces oris* | CHGSTAGEFGNDLR |
| *Lysinibacillus contaminans* | CTEDGEQR |
| *Lysinibacillus sinduriensis* | CAEEGTKR |
| *Caryophanon tenue* | CAFDGEER |
| *Vagococcus lutrae* | CDKPNYTEKR |
| *Carnobacterium divergens* | CVGEVGTVWR |
| *Pediococcus inopinatus* | CNWTGSMR |
| *Aerococcus christensenii* | CGSFNDTSER |

| | |
|---|---|
| *Aerococcus sp. HMSC062B07* | CASFDDISER |
| *Macrococcus caseolyticus* | CTDIKGTAR |
| *Pediococcus ethanolidurans* | CASPYTNEPNR |
| *Nosocomiicoccus sp. HMSC09A07* | CGTLDGASR |
| *Nosocomiicoccus sp. HMSC059G07* | CGTLDGVNR |
| *Pediococcus ethanolidurans* | CDDTGTGR |
| *Massilibacterium senegalense* | CDVPTPTKNR |
| *Finegoldia magna* | CDMPNDPDNR |
| *Catellicoccus marimammalium* | CTLDHDNSGKTLYR |
| *Peptostreptococcus russellii* | CYKDEEGYR |
| *Peptostreptococcus anaerobius* | CYYDDSNYR |
| *Peptostreptococcus sp. MV1* | CYYDVEGYR |
| *Peptostreptococcus sp. D1* | CYYDEAGYR |
| *Catellicoccus marimammalium* | CNTSGDQR |
| *Exiguobacterium profundum* | CTFDTTER |
| *Exiguobacterium sp. AT1b* | CTFDTTER |
| *Exiguobacterium chiriqhucha* | CTFDATER |
| *Exiguobacterium sp. ZOR0005* | CTFDATER |
| *Exiguobacterium sp. SH31* | CTFDATER |
| *Alkalibacillus haloalkaliphilus* | CDIPSKPHNR |
| *Virgibacillus proomii* | CDIPSEPFNR |
| *Fructobacillus pseudoficulneus* | CLFPDTTKR |
| *Leuconostoc carnosum* | CLFPSTQYR |
| *Leuconostoc gelidum subsp. gasicomitatum* | CLFPSTQYR |
| *Leuconostoc sp. BM2* | CLFPSTQYR |
| *Leuconostoc mesenteroides subsp.* | CLFPSTAYR |
| *Leuconostoc lactis* | CLFPSTQYR |
| *Leuconostoc citreum* | CLFPSTNYR |
| *Lactococcus sp. RsY01* | CSDVVGEKR |
| *Vagococcus penaei* | CPTPQVTSQR |
| *Pilibacter termitis* | CEGNIGTIYR |
| *Leuconostoc gelidum* | CDEDADFQAHLRVTNYTDFTCNKR |
| *Leuconostoc citreum* | CLEDDAFWAQVKRSGYTNFKADKR |
| *Leuconostoc mesenteroides* | CLEDQEFWQQVKASHYTNFTAKKR |
| *Leuconostoc lactis* | CLEDADFWRQVKASGYTNFHAPKR |
| *Leuconostoc gelidum* | CLEDAAFWQEVKASHYTNFHADKR |
| *Leuconostoc gelidum* | CDEPGWFNLHPENR |

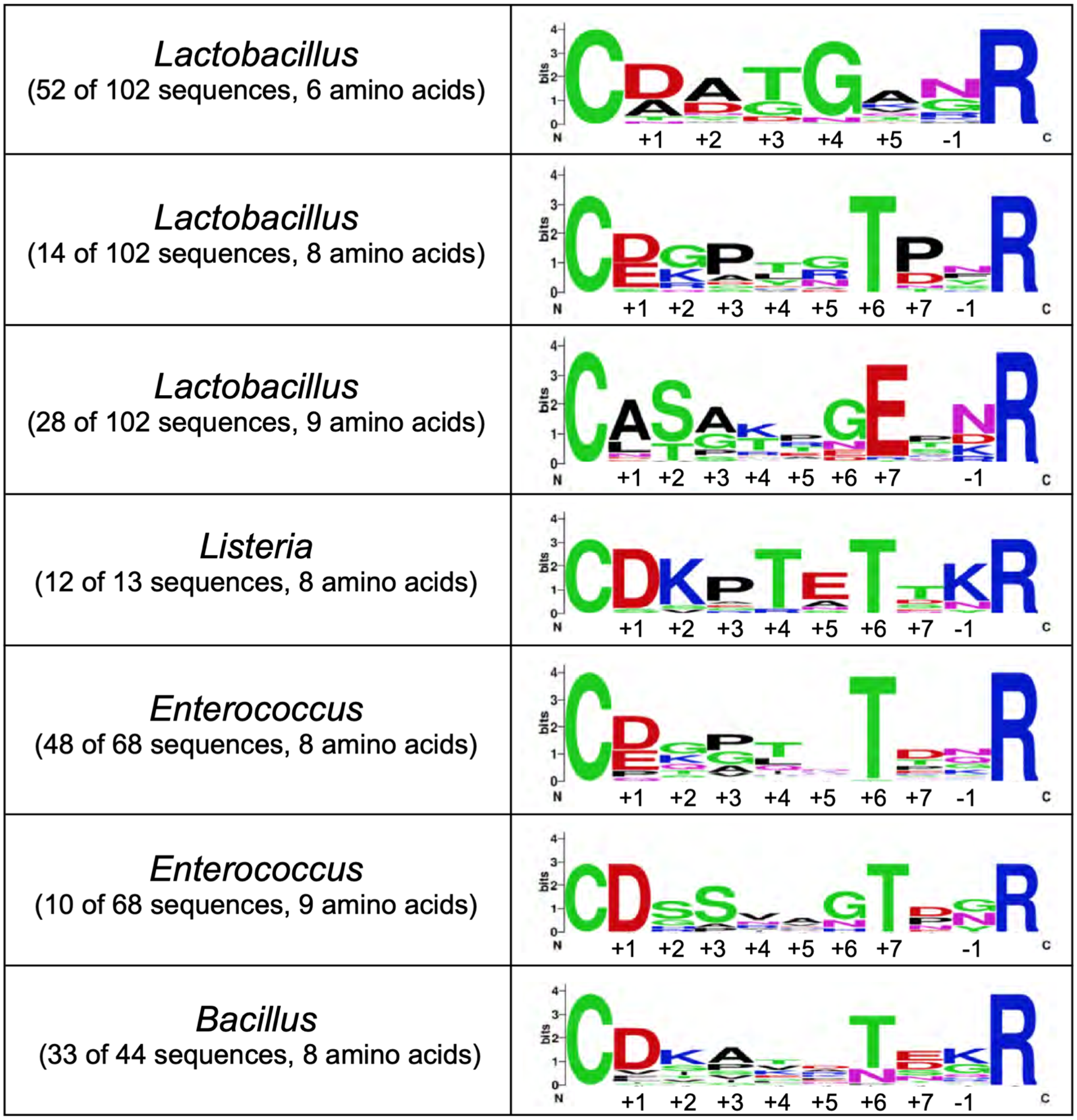


**Fig. S8. WebLogo analyses of β7-β8 loop sequences in multiple genera.** WebLogo analyses of β7-β8 loop sequences of the same lengths are shown for multiple genera, including *Lactobacillus* (loop lengths of 6 amino acids, as well as 8 and 9 amino acids), *Listeria* (8 amino acids), *Enterococcus* (8 and 9 amino acids), and *Bacillus* (8 amino acids). The number of sequences included in each is under the genera name, to the left of the WebLogo, and all β7-β8 loop sequences are in **Table S3**. Numbering, to the β7-β8$^{+7}$ position (here, as "+7"), and including the β7-β8$^{-1}$ (here, as "-1") is under each sequence.

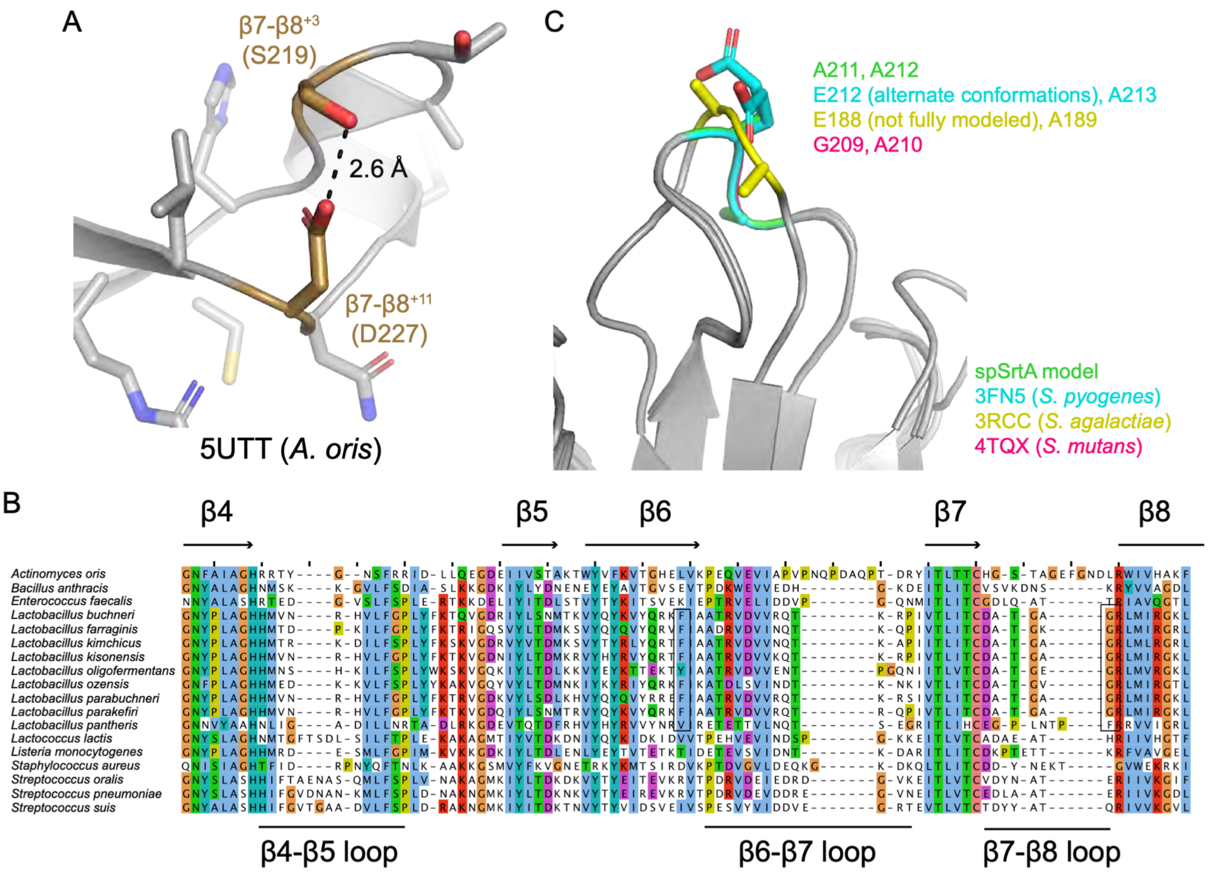


**Fig S9. Sequence analysis of several SrtA proteins supports predicted β7-β8 interactions.** (A) The structure of *A. oris* SrtA (PDB ID 5UTT) shows a hydrogen bond formed between β7-β8$^{+3}$ S219 and β7-β8$^{+11}$ D227 (colored gold, with side chain sticks colored by heteroatom). The distance is 2.6 Å, as labeled. The rest of the protein is in gray cartoon with side chains in stick representation and colored by heteroatom (N=blue, O=red). (B) The sequence alignment of several SrtA proteins that do not contain a charged or polar residue at β7-β8$^{-1}$ show that these proteins all contain a hydrophobic residue that can favorably interact with a hydrophobic residue at the β6$^{-2}$ position. (C) *Streptococcus* structures are shown in gray cartoon, with the midpoint residue of the β7-β8 loop shown in stick representation and colored as labeled (with O=red and N=blue). These residues are all exposed to solvent and do not appear to make specific intra-protein interactions.

### Supplemental Experimental Procedures for Peptide Synthesis

*General Synthetic Procedures.* All peptide substrates were synthesized via manual Fmoc solid phase peptide synthesis (SPPS) using Fmoc Rink amide MBHA resin (synthesis of individual sequences) or SynPhase lanterns (tandem synthesis of multiple sequences) as the solid support. All steps (washing, coupling, deprotection) were performed at room temperature and included gentle agitation on a bench-top rocking platform. All materials, including standard Fmoc amino acids, Fmoc Rink amide MBHA resin, Mimotope SynPhase lanterns (PSLRAM015), and reagents for coupling, deprotection, and resin cleavage were obtained from commercial sources and used without further purification. Incorporation of the 2,4-dinitrophenyl (Dnp) chromophore was achieved using a commercially available lysine building block (Fmoc-L-Lys(Dnp)-OH) purchased from ApexBio. Boc-2-aminobenzoic acid was obtained from Chem-Impex International. A colorimetric ninhydrin test kit for monitoring coupling reactions was purchased from Anaspec.

*Synthesis of Individual Sequences.* Peptides that were independently synthesized utilized Fmoc Rink amide MBHA resin as the solid support. First, a 15 mL polypropylene synthesis vessel fitted with appropriate frits and inlet/outlet caps was loaded with Rink resin at a 0.1 mmol scale. The resin was swollen prior to synthesis with ~10 mL N-methyl-2-pyrrolidone (NMP) (3x, 10 min per wash). Following the NMP washes, the base-labile Fmoc protecting group was removed with 20% piperidine in NMP (2x, 10-20 min), followed by washing with ~10 mL of NMP (3x, 5 min per wash). For each added residue, the following coupling solution was prepared in a 3 mL glass vial: Fmoc amino acid (0.3 mmol, 3.0 equivalents relative to resin loading), O-(benzotriazol-1-yl)-N,N,N',N'-tetramethyluronium (HBTU) (0.3 mmol), N-N-diisopropylethylamine (DIPEA) (0.5

mmol) dissolved in 3 mL of NMP. Following thorough mixing, this solution was added to the synthesis vial with the deprotected resin along with ~3 mL of additional NMP to fully suspend the resin. Couplings were incubated for a minimum of 1 hour at room temperature. Following each coupling, the resin was washed with ~10 mL NMP (3x, 10 min per rinse). The resin was then deprotected with ~10 mL of 20% piperidine in NMP (2x, 10-20 min per treatment), and washed with ~10 mL NMP (3x, 5 min per wash). Repeated cycles of coupling and deprotection were then used to assemble the target sequence. An aminobenzoyl (Abz) fluorophore was installed at the N terminus of all peptides through the coupling of Boc-2-aminobenzoic acid using the same coupling protocol described above. Following the synthesis of the desired sequence, the resin was washed with ~10 mL of NMP (3x, 10 min per wash), followed by ~10 mL of dichloromethane (DCM) (3x, 10 min per wash). A 5 mL solution of 95:2.5:2.5 TFA/TIPS/$H_2O$ was used to cleave most peptide from the resin (2x, 30 min per treatment). Peptides containing cysteine or methionine were cleaved using a solution of 90:2.5:2.5:5 TFA/TIPS/EDT/thioanisole. Peptides containing tryptophan required a cleavage solution of 88:5:5:2 TFA/phenol/$H_2O$/TIPS. The cleaved peptide solutions were concentrated via a rotary evaporator, and the remaining residue was added dropwise to 35 mL of diethyl ether chilled over dry ice. The suspension was then centrifuged at 4500 rpm for 5 minutes at $4^o$ C to collect the precipitated peptide. The diethyl ether was decanted and the crude peptide was dried overnight under vacuum.

*Tandem Peptide Synthesis.* SynPhase lanterns (0.015 mmol loading capacity) were used in order to discretely synthesize numerous peptides in tandem. For parallel coupling of the same residue, multiple lanterns were loaded into a single 15 mL polypropylene synthesis vessel, and deprotection and rinsing were carried out in the same manner and volume as described above for individual peptide sequences. Coupling solutions for attaching Fmoc amino acids were also prepared similarly, with at least a 3x molar excess of the Fmoc amino acid and HBTU, and at least a 5x molar excess of DIPEA. The volume of NMP was adjusted according to the number of lanterns used in order to maintain reagent concentrations consistent with those used in the synthesis of individual peptide sequences. In order to couple the residues that varied between the peptides, the lanterns were transferred from the 15 mL synthesis vessel to individual 3 mL glass vials containing 1 mL of the appropriate coupling solution. Couplings were incubated for at least 2 hours and were not agitated. Prior to being returned to the synthesis vessel, the lanterns were washed with ~3 mL of NMP (3x, 5 min per wash) to prevent cross-contamination of the coupling solution. A fourth wash with ~10 mL NMP was carried out once the lanterns were transferred back to the original synthesis vessel. Once the peptides were complete, each lantern was moved to individual 3 mL vials for cleavage with 1 mL of 95:2.5:2.5 TFA/TIPS/$H_2O$ (2x, 30 min per treatment). Peptides containing cysteine or methionine were cleaved using a solution of 90:2.5:2.5:5 TFA/TIPS/EDT/thioanisole. The cleaved peptide solution was then concentrated using a rotary evaporator. The remaining residue was then added dropwise to a 15 mL polypropylene centrifuge tube containing 2 mL of dry ice-chilled diethyl ether. The majority of sequences precipitated under these conditions and were recovered via centrifugation (4500 rpm for 10-20 minutes at 4 $^{o}$C). The diethyl ether was then decanted and the crude peptides were dried overnight under vacuum. In cases where the peptides were not effectively precipitated from ether, they were suspended in ~10 mL of water and lyophilized.

*Peptide Purification.* Crude peptides from both independent and tandem synthesis were resolvated in a minimum volume of MeCN and $H_2O$ and were purified via RP-HPLC [Phenomenex Luna 5 μm, 100 Å C18 column (10 x 250 mm), aqueous (95:5 $H_2O$/MeCN, 0.1% formic acid) / MeCN (0.1% formic acid) mobile phase at 4.0 mL/min, method: hold 20% MeCN (0.0-2.0 min), linear gradient of 20-90% MeCN 2.0-15.0 min, hold 90% MeCN 15.0-17.0 min)]. The purified peptide fractions were concentrated via a rotary evaporator and then lyophilized. The identity of each peptide was confirmed via ESI-MS (**Table S3**), and the purity of each peptide was confirmed to be >90% by RP-HPLC.

*Peptide Stock Solution Preparation.* Prior to use in sortase-catalyzed reactions, purified peptides were dissolved in a minimum volume of 10:90 DMSO/$H_2O$, 1:1 DMSO/$H_2O$, or pure DMSO. Peptide concentrations were estimated using the absorbance of the Dnp chromophore at 360 nm (extinction coefficient = 17,300 $M^{-1}cm^{-1}$) (2, 3). The stocks were then diluted to a working concentration of either 1 mM, 5 mM, or 10 mM depending on DMSO content in order to ensure that the final sortase-catalyzed reaction mixtures contained ≤5% DMSO by volume. Specifically, reactions utilizing the Abz-LPATFG-K(Dnp) substrate contained a final DMSO concentration of 5% (v/v), whereas reactions with the remaining peptides substrates contained ~0.5-1.5% DMSO (v/v).
